# Exploring the mycovirome: novel and diverse mycoviruses in *Botrytis cinerea*

**DOI:** 10.1101/2025.03.28.645015

**Authors:** Sarah C. Drury, Abdonaser Poursalavati, Pierre Lemoyne, Dong Xu, Peter Moffett, Odile Carisse, Herve van der Heyden, Mamadou L. Fall

## Abstract

*Botrytis cinerea* is a necrotrophic fungal pathogen that causes significant economic losses to many crops, including vegetables, fruits, and ornamental plants. The management of *B. cinerea* is difficult due to a rise in fungicide resistance. Harnessing mycoviruses that cause reduced virulence (hypovirulence) in *B. cinerea* is a promising alternative. Over 100 mycoviruses have been identified in *Botrytis* spp. to date, including several hypovirulence-inducing mycoviruses. This research aimed to further explore, for the first time in Canada, the mycovirome of *B. cinerea* and identify potential hypovirulence-inducing mycoviruses. Isolates of *B. cinerea* were collected from fruits and vegetables in the province of Quebec. Fitness and pathogenicity criteria, including sclerotia production, colony morphotype, and lesion size were evaluated. A double-stranded RNA (dsRNA) extraction protocol tailored to the detection of mycoviruses was used to sequence dsRNA from 45 isolates with low fitness/pathogenicity, and an in-house bioinformatics workflow was used to profile the mycovirome. Mycoviruses were identified in 44/45 isolates. Most of these had positive single-stranded RNA or dsRNA genomes, and a small number had negative single-stranded RNA, single-stranded DNA, or reverse transcriptase RNA genomes. Following deep analysis of RNA-dependent RNA polymerase and replication initiation proteins, a total of 62 unique contigs were identified belonging to new strains of mycovirus species. Furthermore, four putative novel mycovirus species belonging to *Endornaviridae*, *Botybirnaviridae*, *Peribunyaviridae*, and *Bunyavirales* taxa were identified. Several mycovirus species positively and/or negatively co-occurred with *B. cinerea* isolates collected from strawberry or raspberry. This study revealed a high degree of diversity in the mycovirome of *B. cinerea.* Species accumulation curve analysis indicated that, with the number of isolates characterized, we were unable to capture the full extent of expected diversity. Nevertheless, we identified potential hypovirulence-inducing mycoviruses, including Botrytis cinerea mitovirus 1, Botrytis cinerea hypovirus 1, and Botrytis porri botybirnavirus 1. Some of these novel mycoviruses belonged to taxa known to produce viral particles, which can be an interesting feature for their use as biocontrol agents (BCA).

**Importance:** This study provides the first comprehensive profiling of mycoviruses infecting *Botrytis cinerea* in Canada, a significant step in understanding how these viruses can naturally limit crop disease. Due to growing resistance against conventional fungicides, new biological methods to control *B. cinerea* are crucial. By profiling mycoviruses in fungal samples collected in Quebec, we identified several novel viruses that appear to reduce the pathogenicity of *B. cinerea*. These viruses, known as hypovirulence-inducing mycoviruses, could be used to develop biocontrol agents (BCA), offering a more sustainable disease management alternative. Notably, we found virus families with extracellular potential, which may enable easier application as BCAs in agriculture. This research not only broadens the understanding of fungal virology but also holds promise for innovative, eco-friendly approaches to managing *Botrytis cinerea* in Canada.

## Introduction

*Botrytis* spp. are necrotrophic fungal species infecting many important crops, including vegetables, fruits, and ornamental plants [1]. The major species of *Botrytis*, *B. cinerea* Persoon: Fries (teleomorph *Botryotinia fuckeliana* (from Bary) Whetzel), has been considered the second most scientifically and economically important fungal plant pathogen [2], causing losses of $10– 100 billion annually worldwide [3]. *B. cinerea* is an airborne pathogen that infects over 1000 plant species [4], primarily attacking dicotyledonous plants, but also infecting some monocotyledonous plants [5]. Application of synthetic fungicides is still the main management method, but alternative management strategies are needed due to the rise in fungicide resistance [6–8]. Many fungicides registered for *Botrytis-*disease management, such as quinone outside inhibitors, succinate dehydrogenase inhibitors, benzimidazoles, anilinopyrimidines, dicarboximides, and phenylpyrroles, have single-site activity to increase the specificity of their modes of action [9, 10]. These fungicides can lead to several mechanisms that cause resistance, including target site modifications [10]. In addition, in several countries, broad-spectrum fungicides that are not prone to resistance will soon be removed from registration due to their environmental toxicity, which will increase the dependency and use of single-site fungicides [11].

The virulence of *Botrytis* spp. can be affected by viruses, called mycoviruses, that infect and replicate in fungi They are transmitted between cells through a process known as cytoplasmic mixing, which occurs after hyphal anastomosis and heterokaryosis. Additionally, they can be transmitted vertically by spore dispersal [12–14]. Most mycoviruses lack an extracellular phase; however, the single-stranded DNA (ssDNA) mycoviruses, Sclerotinia sclerotiorum hypovirulence-associated DNA virus 1 (SsHADV-1), Fusarium graminearum gemyptripvirus 1 (FgGMTV), and Botrytis gemydayirivirus 1 (BGDaV1) have shown extracellular activity when their viral particles are exogenously applied to their respective hosts, *Sclerotinia sclerotiorum*, *Fusarium graminearum*, and *B. cinerea* [15–17]. Mycoviruses with extracellular activities could be ideal candidates as biocontrol agents (BCA) because they can move easily between cells, overcoming the limitations of vegetative incompatibility in the horizontal transfer of mycoviruses.

While most mycoviruses do not affect fungal physiology, some viruses reduce the virulence of fungi, a phenomenon known as hypovirulence. Conversely, there are also mycoviruses that increase the virulence of fungi, also known as hypervirulence. Mycoviruses that cause hypovirulence may be good candidates as BCA. Symptoms of hypovirulence include suppressed fungal growth, reduced sporulation, changes in colony morphology, and loss of fertility during sexual reproduction. The most widely studied use of hypovirulence-inducing mycoviruses as BCA is in the management of *Cryphonectria parasitica*, the causative agent of chestnut blight, in European chestnut forests. A hypovirulence-inducing mycovirus, Cryphonectria hypovirus 1, was transmitted among *C. parasitica* populations and contributed to a decline of chestnut blight in Europe In addition, in field conditions, spraying a strain of *S. sclerotiorum* infected with SsHADV-1 converted *S. sclerotiorum*, the causative agent of white mold, into a beneficial endophyte in rapeseed, reducing stem rot severity by 67.6% and increasing the yield by 14.9% [21].

Advances in high-throughput sequencing technologies have allowed for the discovery of many mycoviruses, and for mycoviral diversity and evolution to be studied. Surveys of the mycovirome of phytopathogenic fungi such as *S. sclerotiorum*, *B. cinerea*, *Rhizoctonia* spp., *Fusarium* spp., *Colletotrichum truncatum*, *Macrophomina phaseolina*, *Diaporthe longicolla*, *Pyricularia oryzae*, *Ustilaginoidea virens, Rosellinia necatrix*, *Alternaria* spp., and *Monilinia fructiola*, have led to the identification of many novel mycoviruses [23–32]. Most identified mycoviruses have dsRNA or positive single-stranded RNA (ssRNA(+)) genomes, but mycoviruses with negative-sense RNA (ssRNA(−)), reverse transcriptase RNA (RT ssRNA), and single-stranded DNA (ssDNA) genomes have also been identified [33, 34]. Over 100 mycoviruses have been identified in *Botrytis* spp. [30, 35–45]. Double-stranded RNA was detected in 143 of 200 *B. cinerea* isolates collected from strawberry, glasshouse tomatoes, grape, cucumber, kiwifruit, French bean, and blackberry crops in New Zealand [46]. In addition, dsRNA elements were found in 53 of 96 *B. cinerea* isolates collected from cucumber, green bean, pepper, tomato, aubergine, zucchini, and grapevine in Spain [38]. Furthermore, 92 mycoviruses were identified in a metatranscriptomic study focused on evaluating the mycovirome of 248 *B. cinerea* isolates collected from grapevine in Italy and Spain [30]. Similarly to what has been observed in other fungi, the majority of mycoviruses found to infect *B. cinerea* have dsRNA or ssRNA(+) genomes [30, 38]. However, mycoviruses with ssRNA(−) genomes [30, 47, 48] and ssDNA genomes have also been identified [17, 30, 44, 49]. Several hypovirulence-inducing mycoviruses have been reported in *B. cinerea*, including Botrytis cinerea hypovirus 1 (BcHV1), Botrytis cinerea mitovirus 1 (BcMV1), Botrytis cinerea RNA virus 1 (BcRV1), and Botrytis porri botybirnavirus 1 (BpBV1) [17, 35, 37, 40–43].

In this report we have sought to 1) determine the taxonomical profile, incidence and diversity of mycoviruses in *B. cinerea* isolates with abnormal colony morphotypes and/or low virulence collected in Quebec, Canada, and 2) evaluate the effect of the *B. cinerea* host plants on the mycovirome composition and abundance. We used a dsRNA extraction method tailored specifically for mycovirus detection and characterization which produced high yields and consistent results. Viral metagenomics was then conducted on 45 isolates of *B. cinerea* with abnormal colony morphotypes and/or low virulence to characterize the mycovirome. A total of 72 previously identified mycovirus species belonging to 20 families as well as several taxonomic ranks that are unclassified at the family level were identified. This included several mycoviruses previously reported to have hypovirulence-inducing effects. Further, 62 unique (not previously reported) contigs belonging to 44 new mycovirus species strains and four putative novel mycovirus species were identified based on RNA-dependent RNA polymerase (RdRp) and replication initiation protein (Rep) analysis.

## Results

### Botrytis cinerea isolate morphotype/virulence evaluation

A total of 206 *B. cinerea* isolates were collected from raspberry, strawberry, and grapevine plants in Quebec, Canada. To increase the chance of identifying hypovirulence-inducing mycoviruses, 42 isolates were selected for sequencing based on producing small or no lesions on bean leaves and/or failure to form or development of small sclerotia (Supplementary Table 1). Three more virulent isolates were included to provide a basis for comparison. For colony morphotype, failure to develop conidia and/or sclerotia or production of small sclerotia can be due to hypovirulence-inducing mycoviruses [35, 37, 40, 50, 51]. Of the 45 selected isolates, 13% had a mycelial morphotype and did not form sclerotia as these isolates had M1, M2, and M3morphotypes (Fig. 1A,B). In addition, 20% of isolates produced small sclerotia, having an S4 or S4/M2 morphotype. A further one isolate (Bc2021-12) only produced sclerotia around the edge of the Petri dish (S1 morphotype). One isolate (Bc2020-236) did not grow so did not produce a morphotype. The remaining isolates produced large sclerotia: 42%, 16%, and 4% had S3, S2, and S3/M3, morphotypes, respectively (Fig. 1A,B). No isolate had a M4 morphotype. Isolate Bc2019-174 has regular virulence and the lesion size caused by this isolate was used as a reference for the detected of isolates considered to be hypovirulence-inducing. Bc2019-174 caused average lesion sizes of 756 mm^2^ and presented an M2 morphotype on plates. In total, 42 isolates produced reduced lesion sizes compared to Bc2019-174; their lesion sizes were under 440 mm^2^ and four isolates produced no lesion (Fig. 1C-D). Two isolates, (Bc2020-5 and Bc2021-9) were selected as being higher-performing isolates as they produced similar lesion sizes to Bc2019-174 (Fig. 1D).

**Fig 1.**
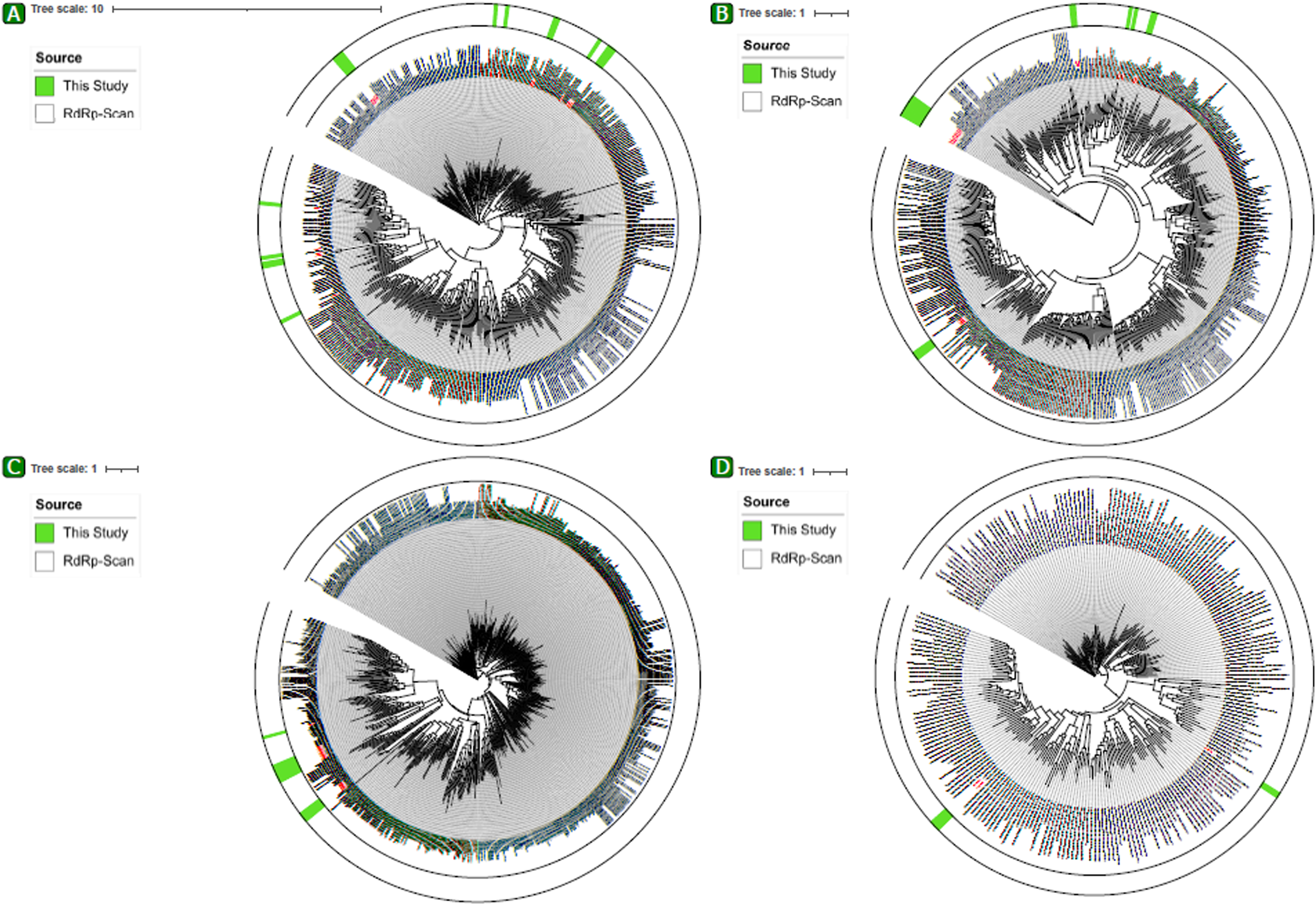
Colony morphology and lesion sizes of *Botrytis cinerea* isolates. **(A)** Isolates were grown on potato dextrose agar (PDA) Petri plates in the dark for three weeks and classified for morphotypes. Images of representative morphotypes from each class are shown, as indicated. Pie chart showing the isolates belonging to different colony morphotypes **(C)** Different lesion sizes determined on detached bean leaves (20°C, 12-h photoperiod, 72 h) **(D)** Average lesion sizes of three replicates of isolates on bean leaves after 72 h were compared to those induced by Bc2019-174 (in orange), a control isolate causing ‘normal’ lesion sizes. Significant differences based on Dunnett’s test, \**P* ≤ 0.05.

### Virus identification and taxonomy

For each isolate, dsRNA was extracted, which was followed by sequencing with the Illumina MiSeq platform. An in-house pipeline called SOVAP was used to identify mycoviruses. After exclusion of the positive control (PvEV-1), phages, and other obvious non-mycovirus taxa, and prior to the identification of novel contigs, the analysis of sequencing results from the 45 isolates revealed 20 classified families, 21 classified genera, and several taxonomic ranks that are unclassified at the family and genus levels. Viral reads of taxon with ssRNA(+) genomes, such as endornaviruses, fusariviruses, and gammaflexiviruses were predominant, followed by those with dsRNA genomes (Fig. 2A, B). A total of 72 previously identified mycovirus species, with different relative abundances, were detected (Fig. 2C). Mycoviruses were identified in 44/45 isolates; mycoviruses were not found in isolate Bc2020-262. A total of 41 isolates harbored more than one virus, and up to 12 mycoviruses were found in a single isolate (Bc2020-141) (Fig. 2C). Botrytis cinerea endornavirus 2 (BcEV2) had the highest occurrence rate in the isolates, as it was detected in 58% of isolates. This was followed by BpBV1 which was detected in 40% of isolates.

**Fig 2.**
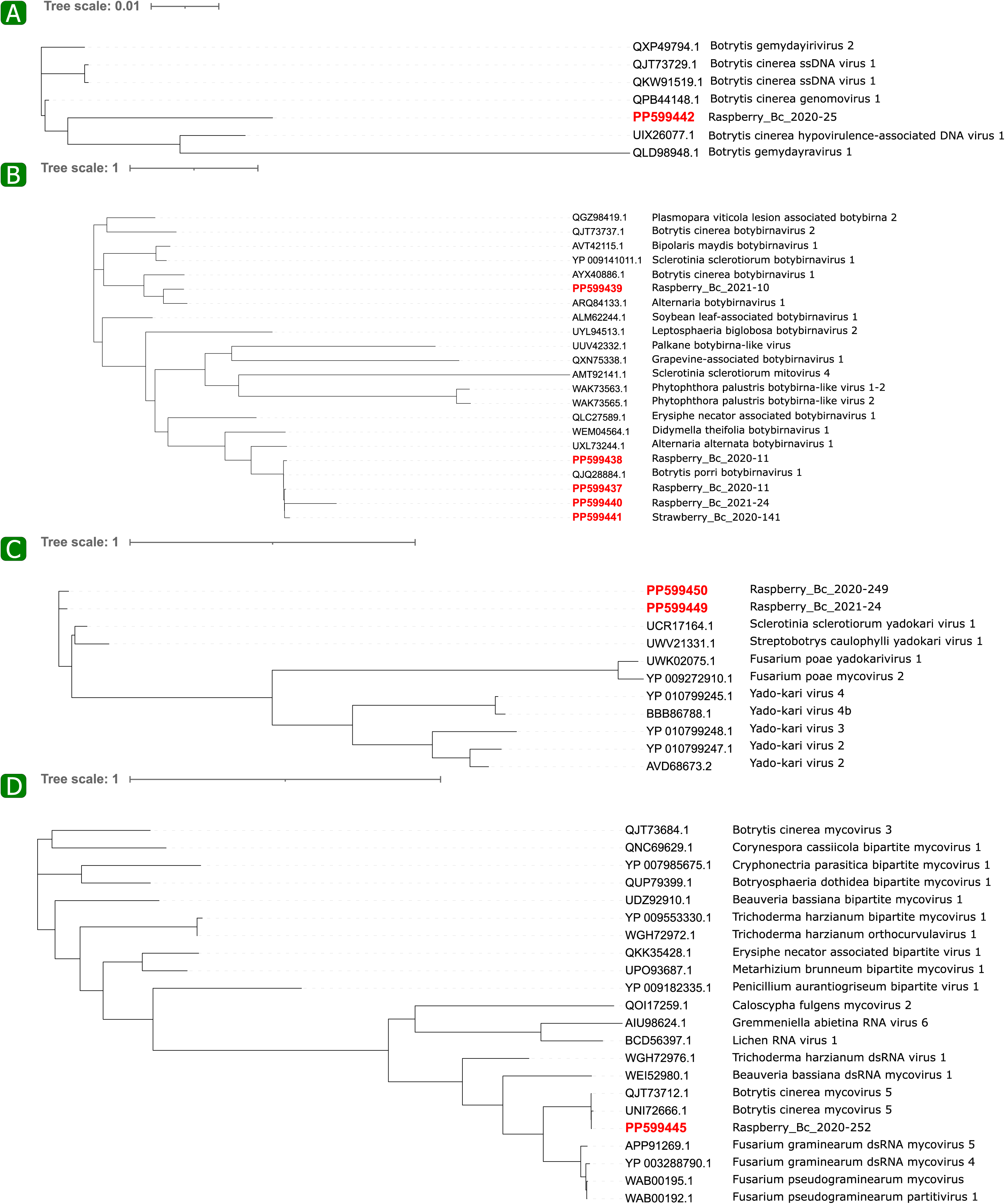
Illustration of the top ten mycovirus taxon at the **(A)** family, and **(B)** genus levels, including groups unclassified at the levels, in *Botrytis cinerea* isolates based on the assigned viral sequences. Stacked bar chart shows the percentage of sequences. **(C)** Heatmap showing the relative abundance (%) of species, based on assigned viral sequences, in *B. cinerea* isolates. The color legend on the right side indicates the relative abundance, ranging from purple (low abundance) to yellow (high abundance).

In addition to the 72 previously identified mycovirus species, 172 unique (not previously reported) viral contigs were initially found using RdRp-Scan based analysis for RNA viruses and manual curation of the sequencing data for DNA viruses. A total of 48 contigs could be assigned to the *Pisuviricota* phylum (Supplementary Fig. 1A), 33 to the *Lenarviricota* phylum (Supplementary Fig. 1B), 38 to the *Kitrinoviricota* phylum (Supplementary Fig. 1C), five to the *Negarnaviricota* phylum (Supplementary Fig. 1D), 24 to the *Botybirnaviridae* family (Supplementary Fig. 1E), seven to the *Cressdnaviricota* phylum, and 17 to taxonomic ranks with ssRNA(+) and dsRNA genomes that are unclassified at the phylum level (Supplementary Fig. 1F).

Following RdRp and Rep motif, and multiple-sequence alignments analyses (Supplementary Fig. 2-4), 66 unique contigs remained – 62 that belonged to 44 new mycovirus species strains and four belonging to putative novel mycovirus species. These contigs were all submitted and validated by an NCBI GenBank team (Table 1). A total of 17 unique contigs remained each for *Lenarviricota* (Fig. 3A; Supplementary Fig. 2A) and *Kitrinoviricota* (Fig. 3B; Supplementary Fig. 2B), 15 remained for *Pisuviricota* (Fig. 3C; Supplementary Fig. 2C), and three unique contigs remained for *Negarnaviricota* (Fig. 3D; Supplementary Fig. 2D). Further, one contig remained in the *Cressdnaviricota* phylum in the *Genomoviridae* family [17] (Fig. 4A; Supplementary Fig. 3A), and five contigs remained for the family *Botybirnaviridae* (Fig. 4B; Supplementary Fig. 3B). Five groups of contigs that are unclassified at the phylum level were also evaluated. Two unique RdRp contigs belonging to the *Yadokariviridae* family were identified based on conserved motifs A-E [53] (Fig. 4C; Supplementary Fig. 3C).

**Table 1.**
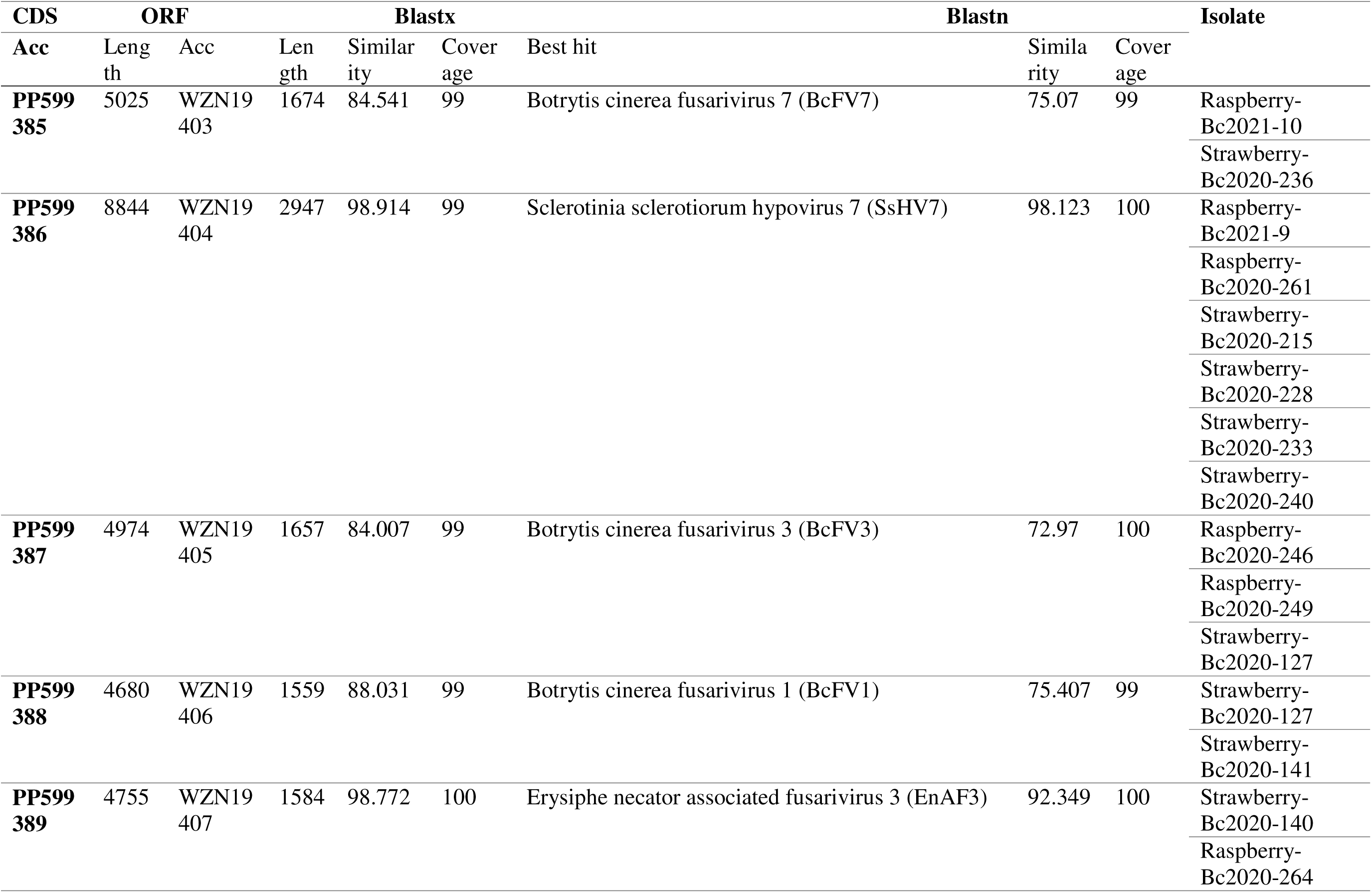
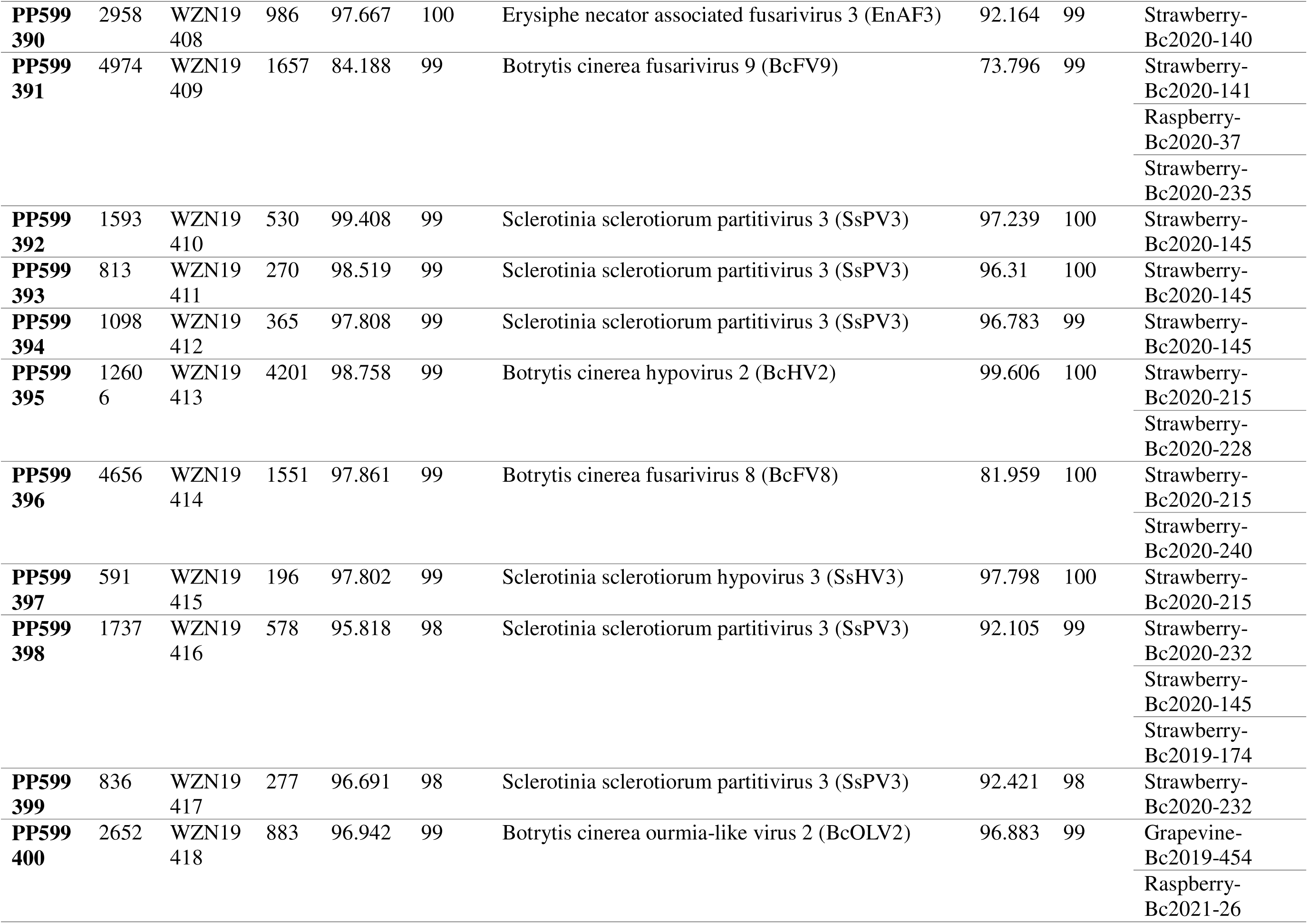
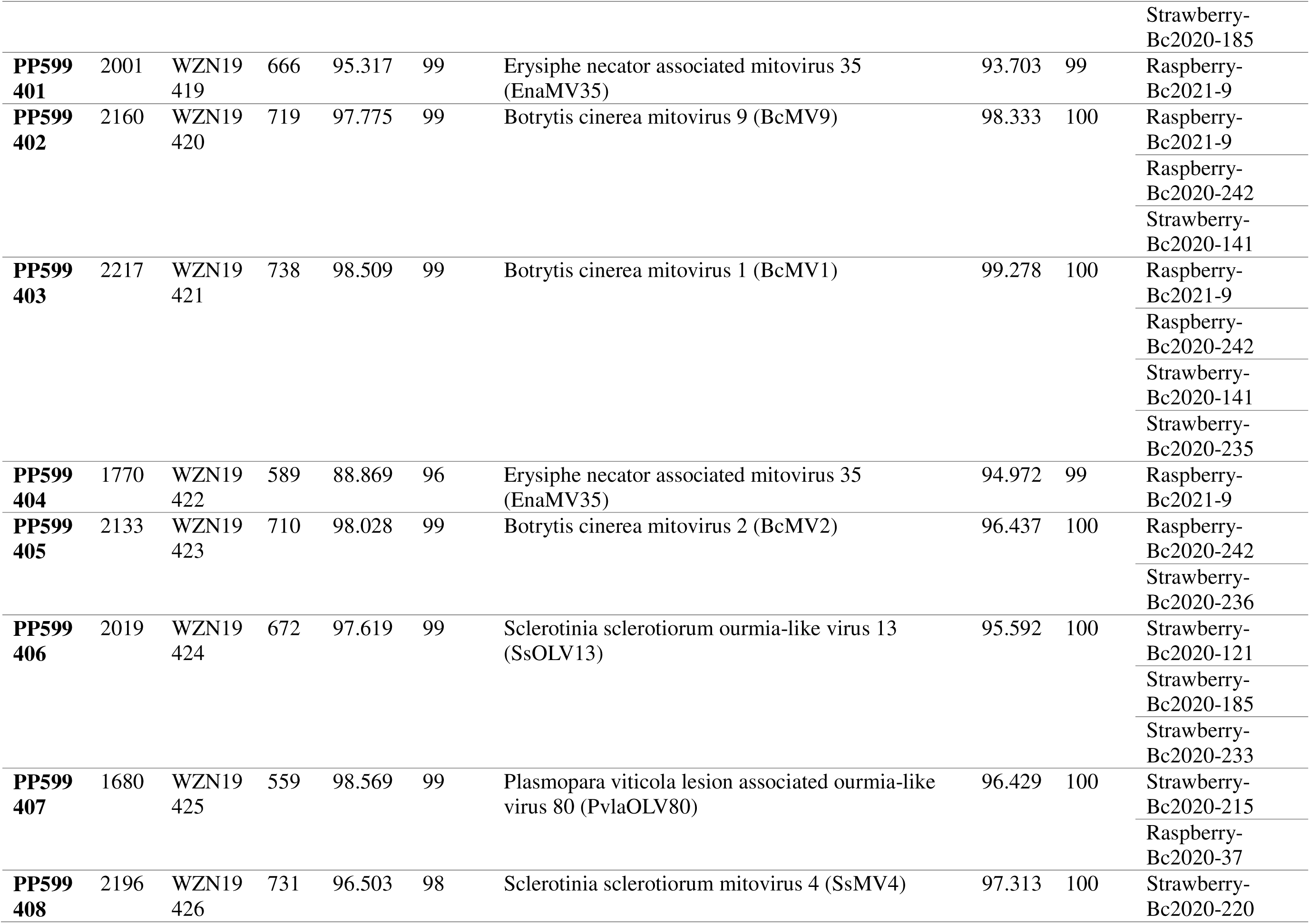
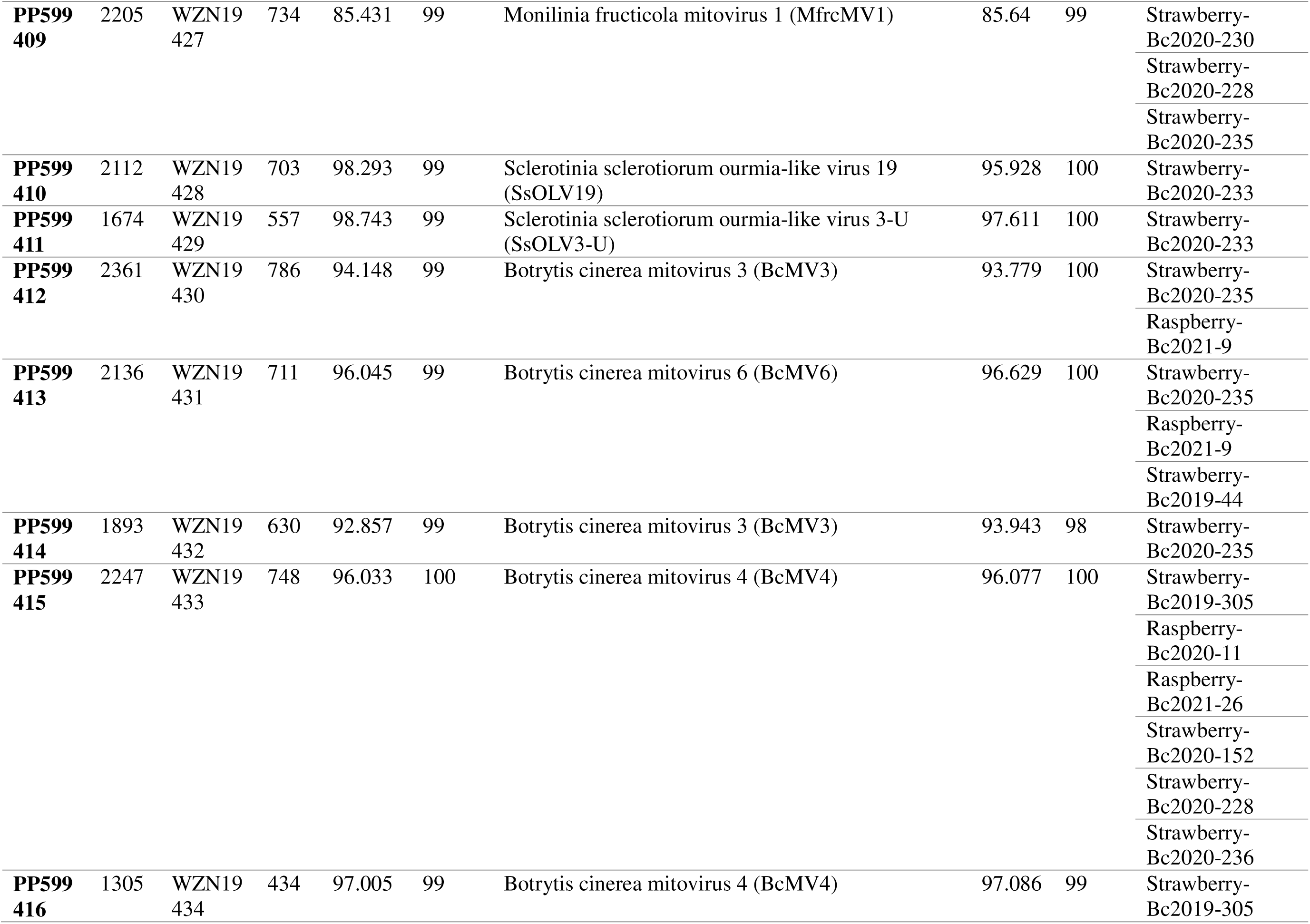
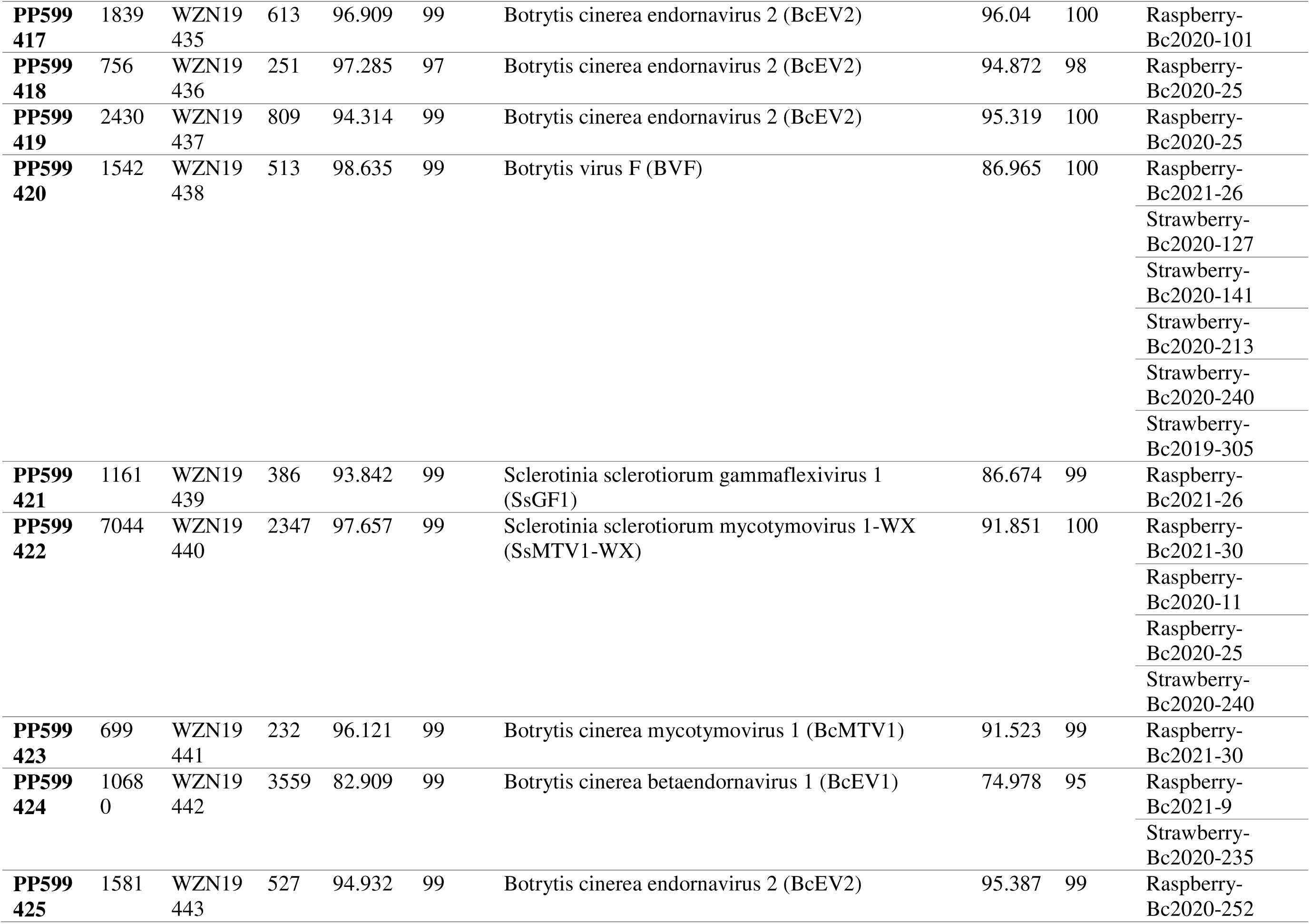
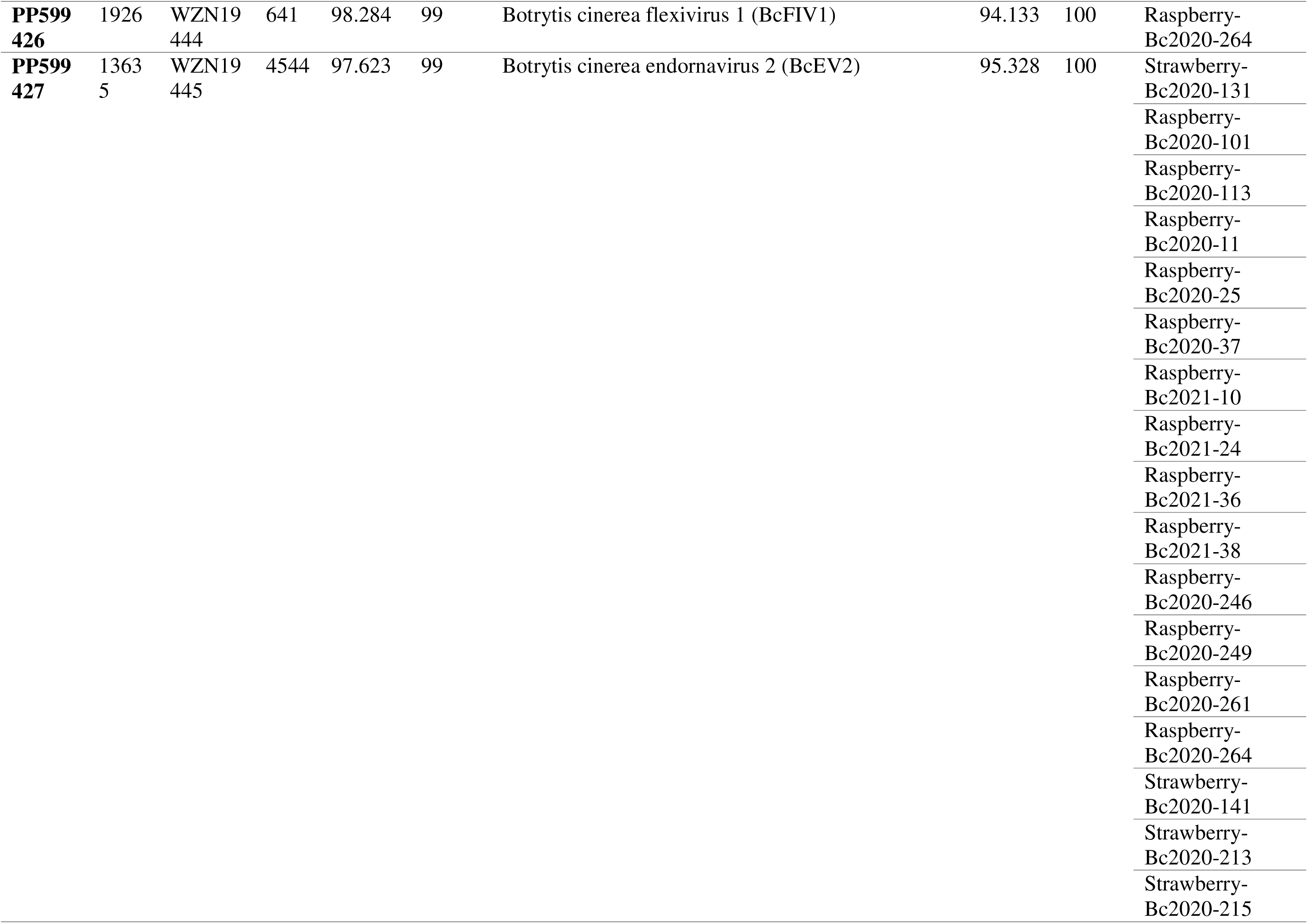
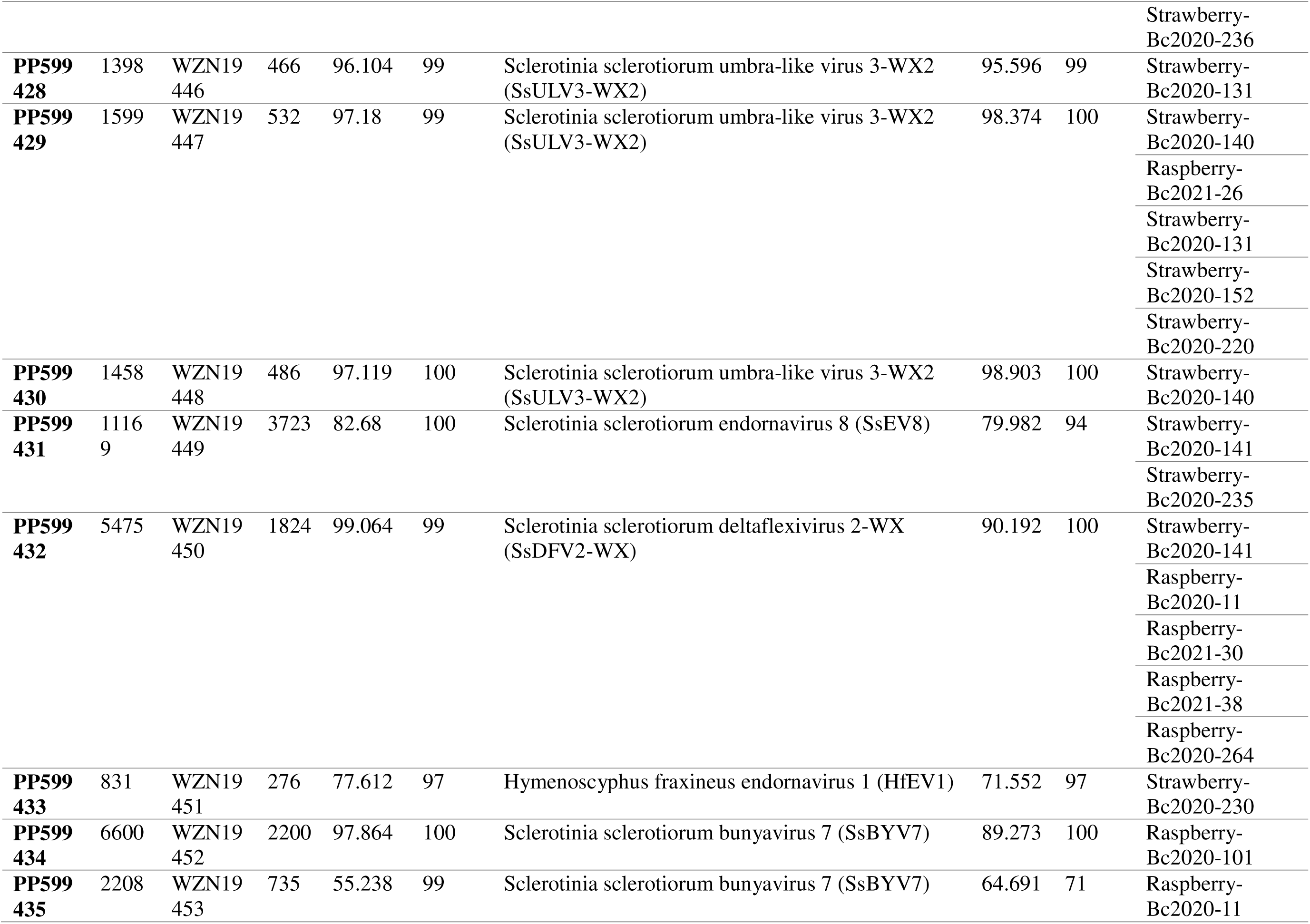
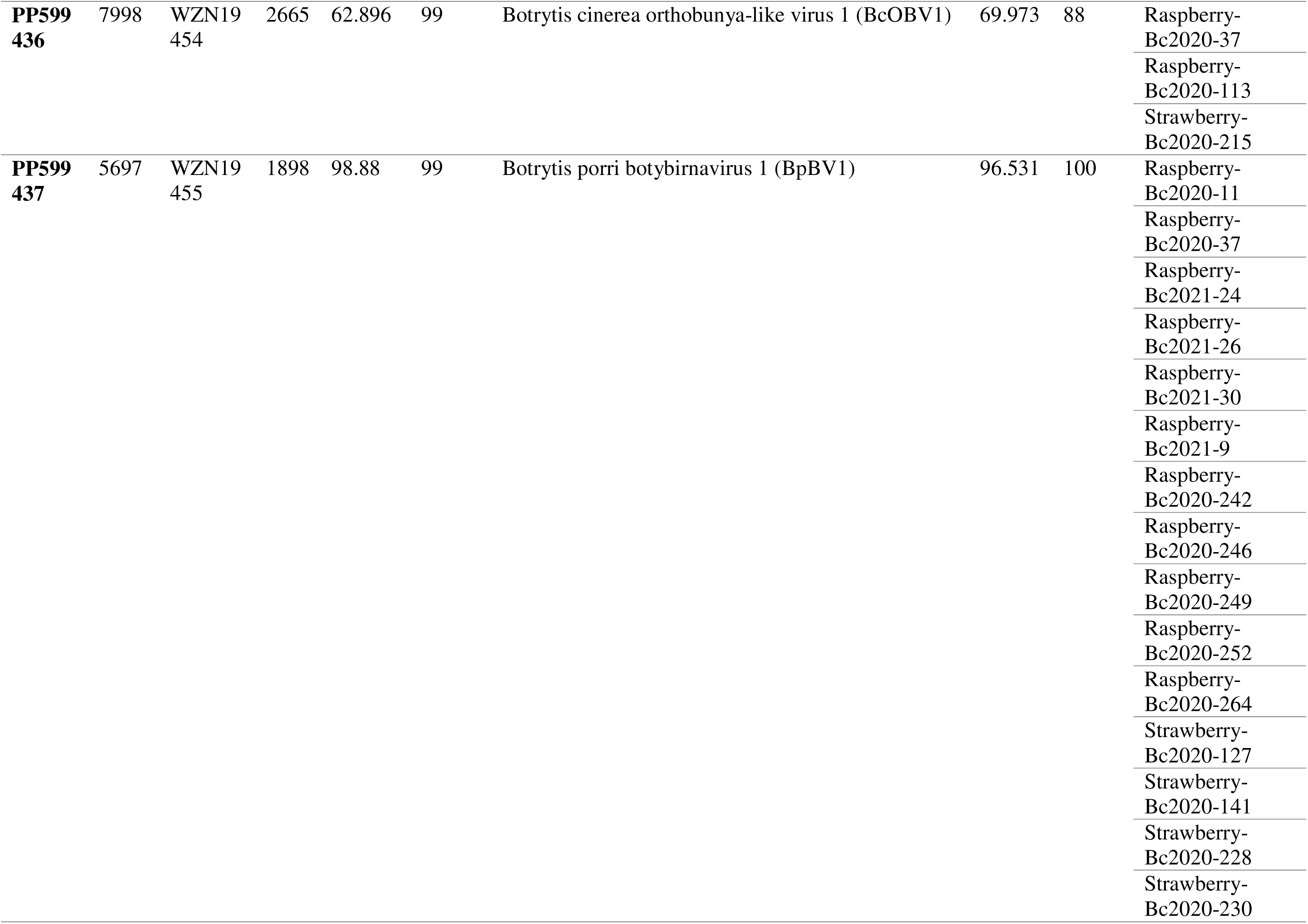
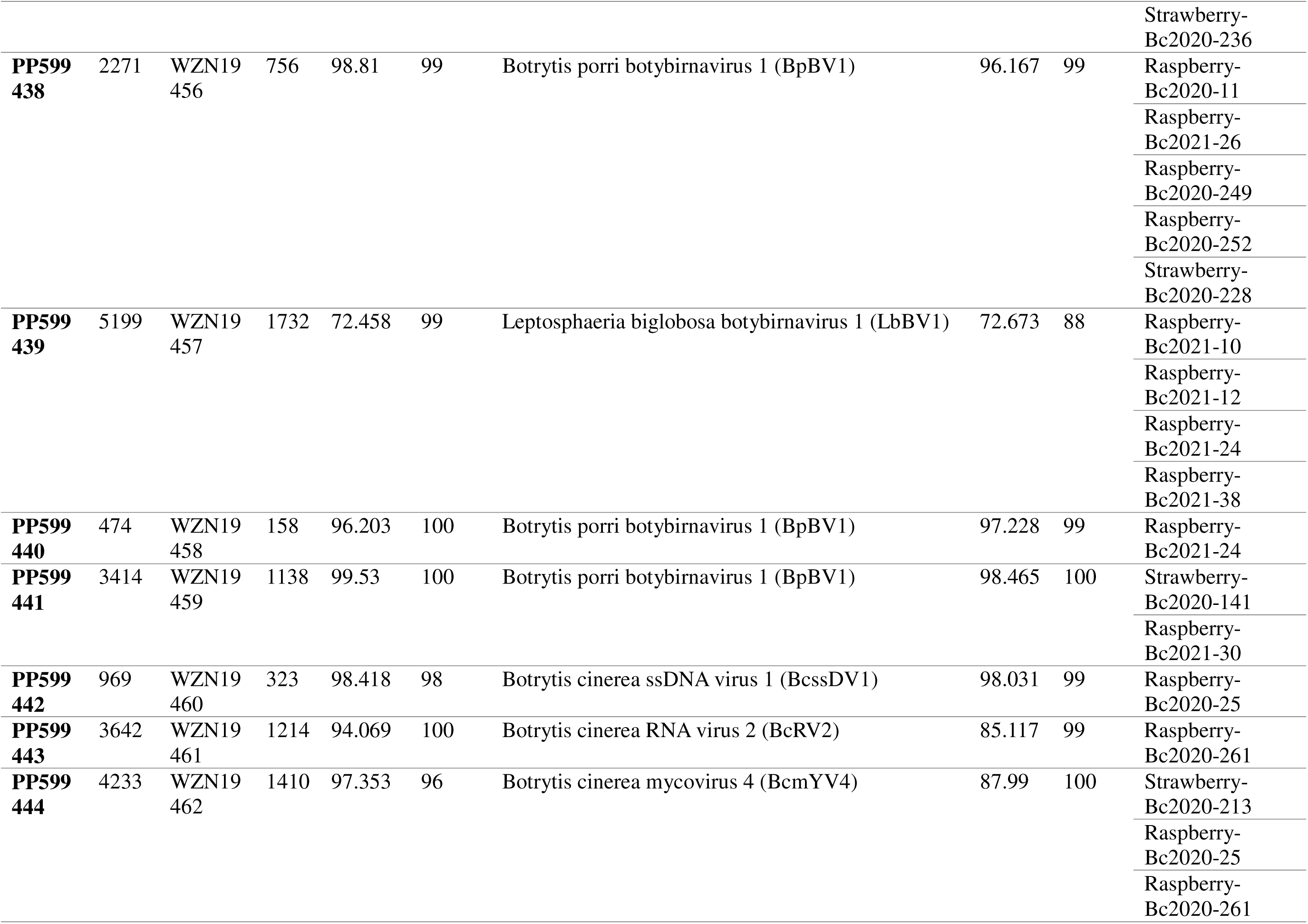
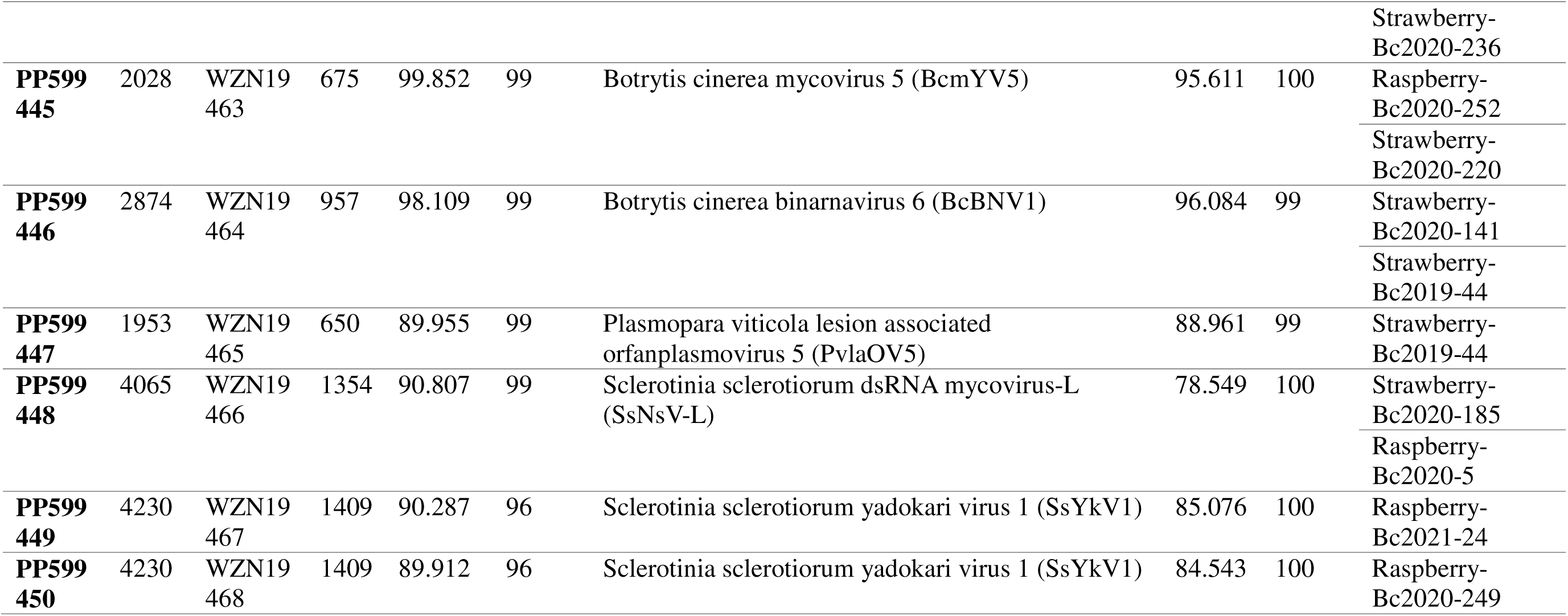
Assembled mycovirus RNA-dependent RNA polymerase (RdRp) contigs with similarity to known viruses identified in isolates of Botrytis cinerea. The isolate with the longest contig candidate from each RdRp cluster is listed first for each contig.

**Fig 3.**
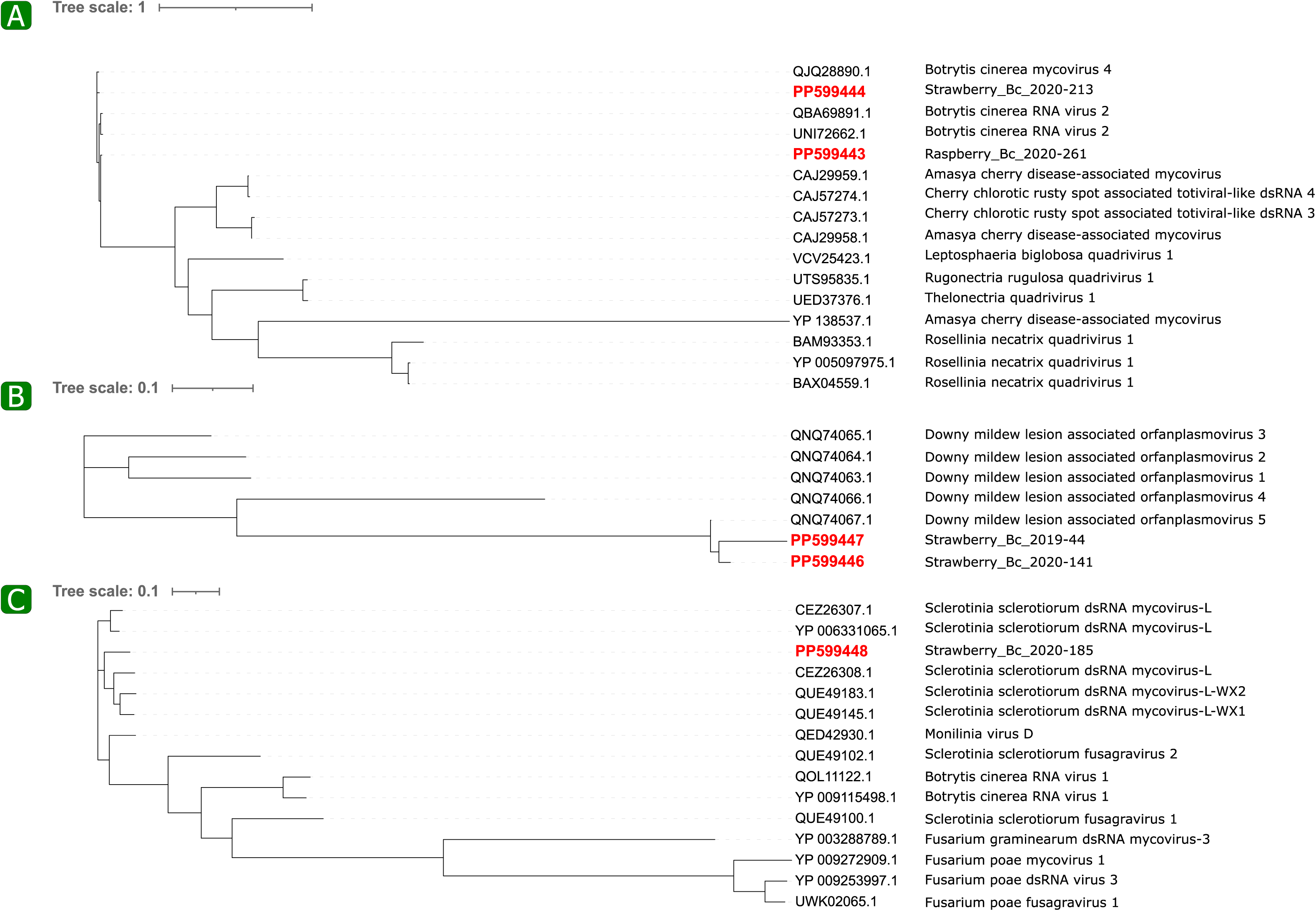
Phylogenetic trees of RNA-dependent RNA polymerase (RdRp) sequences in the **(A)** *Lenarviricota* phylum (n=17), **(B)** *Kitrinoviricota* phylum (n=17), **(C)** *Pisuviricota* phylum (n=15), and **(D)** *Negarnaviricota* phylum (n=5), following protein motif analysis. Accession numbers in red are from this study (indicated in green in the outer ring).

**Fig 4.**
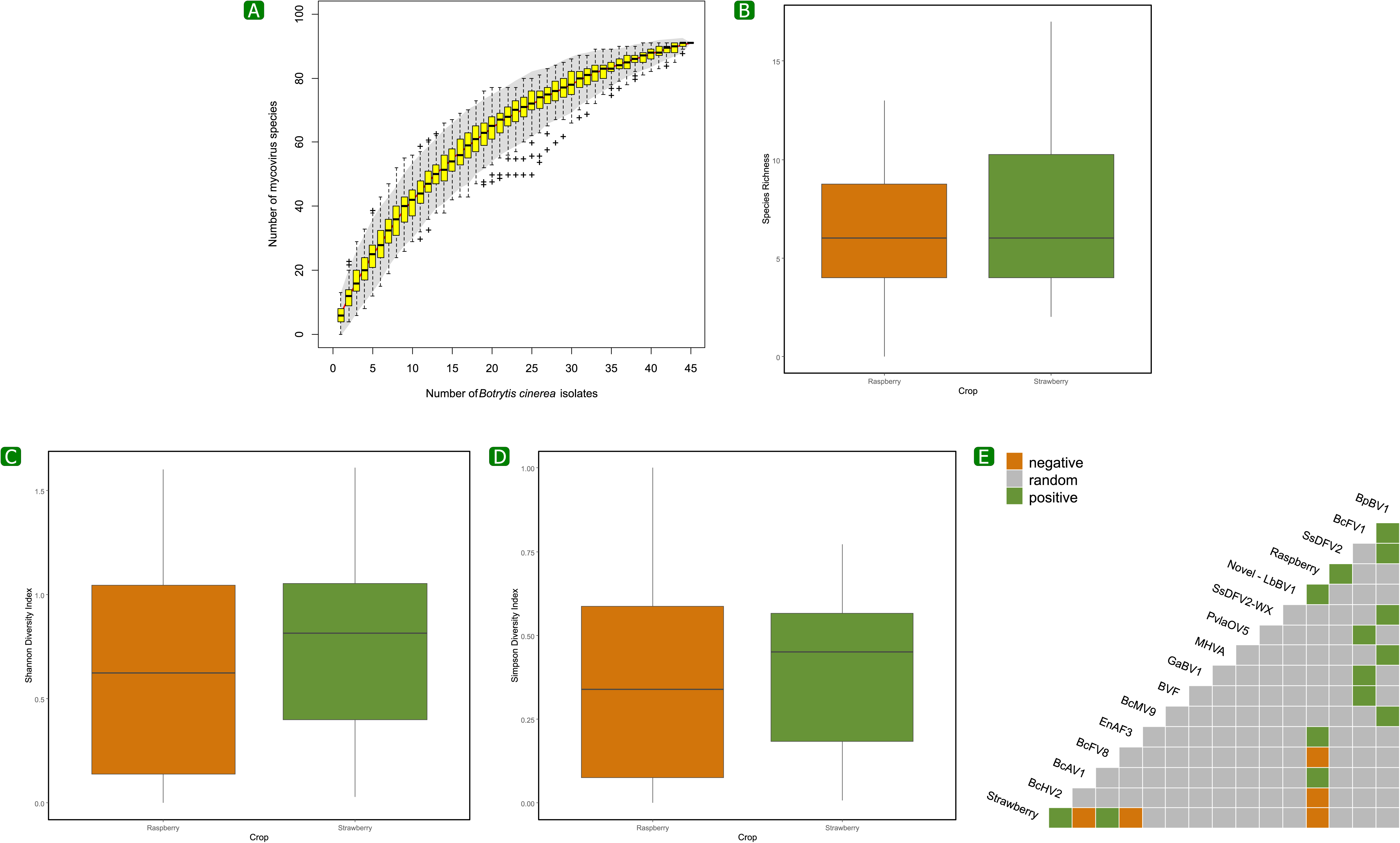
Phylogenetic trees of **(A)** Replication initiation protein sequence in the *Genomoviridae* family (n=1), **(B)** RNA-dependent RNA polymerase (RdRp) sequences in the family *Botybirnaviridae*, (n=5), **(C)** RdRp sequences in the *Yadokariviridae* family (n=2), and **(D)** RdRp sequence related to Botrytis cinerea mycovirus 5 and related dsRNA mycoviruses, following protein motif analyses. Accession numbers in red are from this study (indicated in green in the outer column).

Yadokariviruses are capsidless and are instead trans-encapsidated by an unrelated dsRNA virus in the Duplornaviricota phylum [54]. The three isolates (Bc2020-246, Bc2020-249, and Bc2021-24) that harbored this yadokarivirus also harbored one or more botybirnaviruses which are in the Duplornaviricota phylum. In particular, they all harbored BpBV1, which may be responsible for the trans-encapsidation. One unique RdRp contig was identified that was related to Botrytis cinerea mycovirus 5 and other dsRNA viruses based on analysis of motifs A-E [30] (Fig. 4D; Supplementary Fig. 3D). Two unique RdRp contigs were related to viral RdRps in the *Quadriviridae* family based on motifs I-VIII [30] (Fig. 5A; Supplementary Fig. 4A). Two unique RdRp contigs related to ‘Orfanplasmoviruses’ were evaluated based on domains A-D [55] (Fig. 5B; Supplementary Fig. 4B). Further, a unique RdRp contig was related to Sclerotinia sclerotiorum dsRNA mycovirus L and related dsRNA mycoviruses based on motifs A-E [43] (Fig. 5C; Supplementary Fig. 4C).

**Fig 5.**
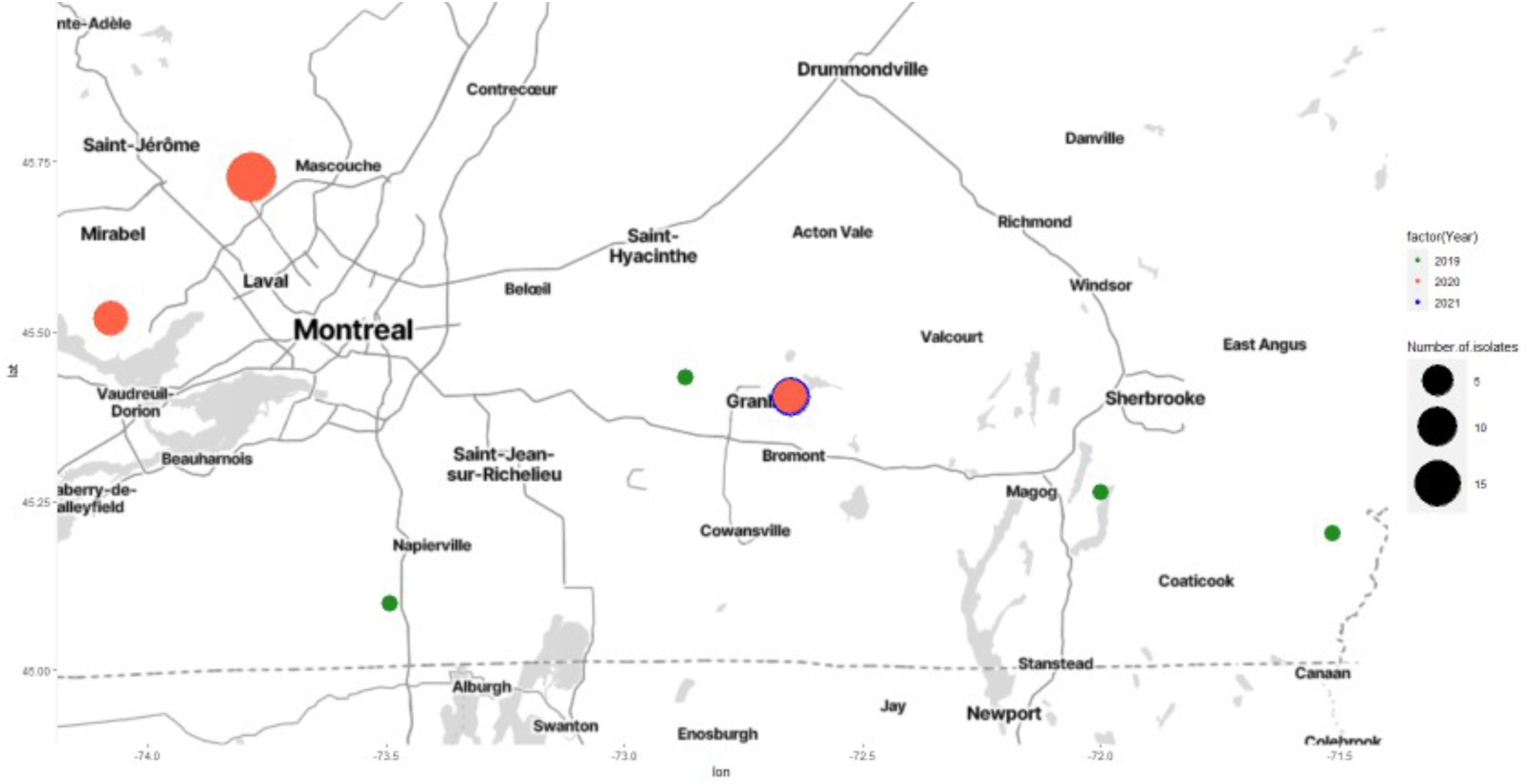
Phylogenetic trees of unique RNA-dependent RNA polymerase (RdRp) sequences **(A)** in the family *Quadriviridae* (n=2), **(B)** related to ‘Orfanplasmoviruses’ (n=2), and **(C)** related to Sclerotinia sclerotiorum dsRNA mycovirus L and related dsRNA mycoviruses (n=1), following protein motif analyses. Accession numbers in red are from this study (indicated in green in the outer column).

The majority of unique contigs identified (62/66) belonged to new strains of previously identified mycovirus species (Table 1). The four remaining contigs belonged to putative novel viral species; a putative novel RdRp viral contig (accession no. PP599433) in the *Kitrinoviricota* phylum was identified in isolate Bc2020-230 in strawberry with 77.6% nucleotide identity to Hymenoscyphus fraxineus endornavirus 1 (HfEV1) based on BlastX (Fig. 3B; Table 1). A putative novel viral contig (accession no. PP599439) belonging to the *Botybirnaviridae* family, found in isolates Bc2021-10, Bc2021-12, Bc2021-24, and Bc2021-38, which were all collected from raspberry, had 72.5% identity to Leptosphaeria biglobosa botybirnavirus 1 (LbBV1) (Fig. 4B; Table 1). Two putative novel viruses were identified in the *Negarnaviricota* phylum. One putative viral RdRp contig (accession no. PP599435) had a 55.2% nucleotide identity to Sclerotinia sclerotiorum bunyavirus 7 (SsBYV7) was identified in Bc2020-11 collected from raspberry (Fig. 3D; Table 1). A second (accession no. PP599436) had a 62.9% nucleotide identity to Botrytis cinerea orthobunya-like virus 1 and was found in Bc2020-37 and Bc2020-113 collected from raspberry, and Bc2020-215 collected from strawberry (Fig. 3D; Table 1).

### Virus diversity and co-occurrence analysis

A total of 81 mycovirus species were identified, when including the novel mycoviruses and new strains of mycoviruses. The majority of mycoviruses had ssRNA(+) genomes (70.3%), followed by dsRNA genomes (23.8%) (Supplementary Fig. 5A). A small number of viruses having ssRNA(−) (4.1%) or ssDNA genomes (1.5%) were identified. The ssDNA viruses, Botrytis cinerea genomovirus 1 (BcGV1), Botrytis cinerea hypovirulence-associated DNA virus 1 (BcHADV1), and Botrytis cinerea ssDNA virus 1 (BcssDV1) were identified, with BcGV1 and BcHADV1 detected in isolate Bc2020-101, and BcssDV1 detected in Bc2020-25. However, the Rep sequences of these three ssDNA viruses share an amino acid identity of 98% and therefore belong to the same species according to Ruiz-Padilla et al. 2023 [50]. Only one virus, Penicillium camemberti virus - GP1, had reverse transcribing (RT) ssRNA genomes (0.4%) (Supplementary Fig. 5A). Mycoviruses identified in this study most commonly belonged to the families *Endornaviridae* (13%), *Botybirnaviridae* (12%), *Fusariviridae* (11%), *Mitoviridae* (10%), and *Hypoviridae* (10%) (Supplementary Fig. 5B).

The full diversity of mycoviruses infecting *B. cinerea* isolates was not captured in our study, as the species accumulation curve did not reach a plateau, suggesting an even greater diversity of mycoviruses in natural populations of *B. cinerea* (Fig. 6A). The species richness and diversity of the mycovirome in the *B. cinerea* isolates collected from strawberries did not significantly (*P* < 0.05) differ from the mycovirome of isolates collected from raspberries. The average viral species richness was not significantly different (*P* < 0.05) in isolates collected from the two crops as it was 6.1 for isolates collected from strawberries and 6.0 for isolates from raspberries (Fig. 6B). A total of 61 mycovirus species were identified both in the isolates from raspberries and from strawberries with 41 species in common between isolates from the two crops. Furthermore, Shannon diversity index and Simpson diversity index did not differ between isolates from strawberry and raspberry (Fig. 6.C-D.)

**Fig 6.**
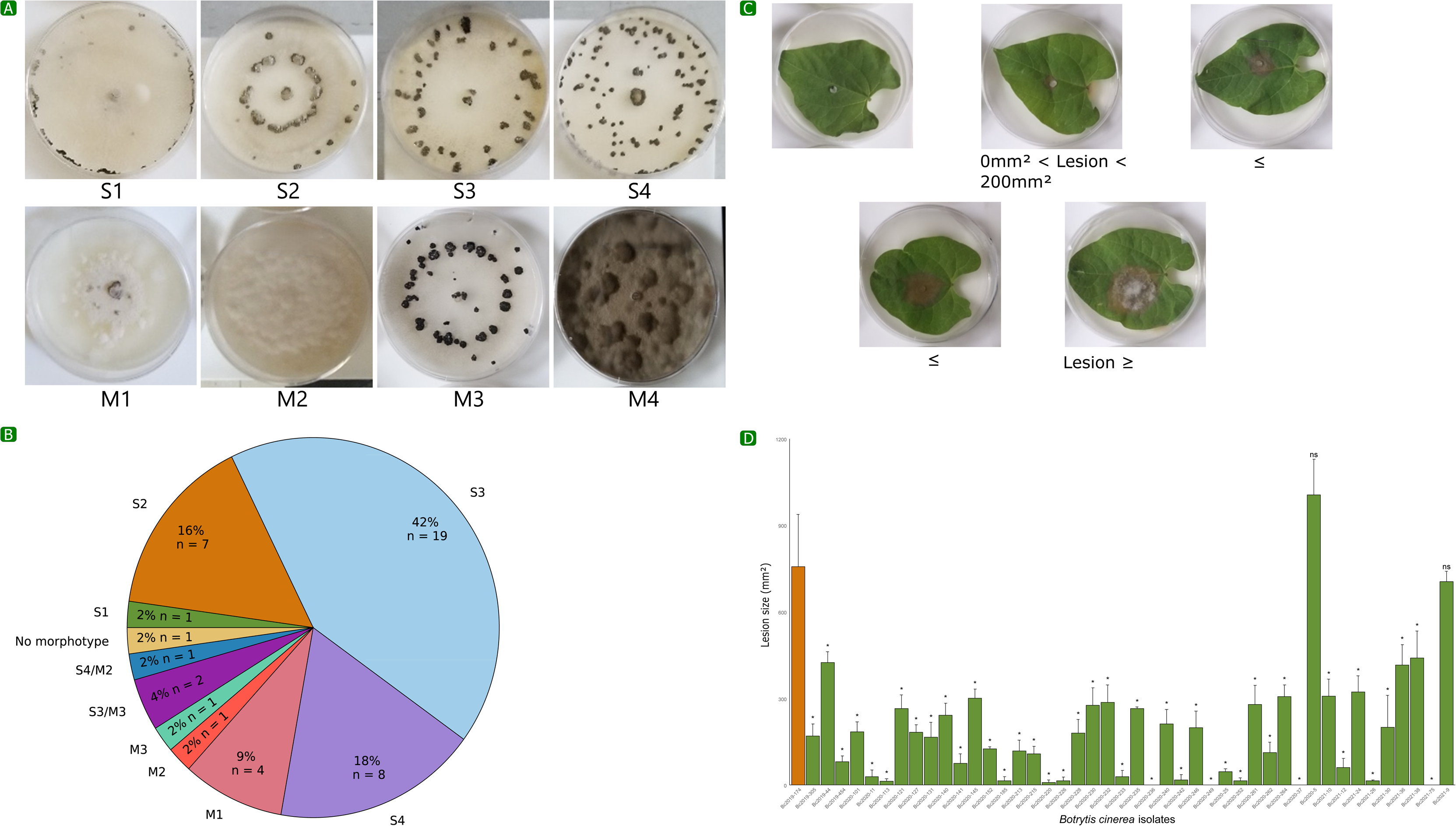
(A) Species accumulation curve of the mycoviruses in *Botrytis cinerea* isolates. The red curve represents the number of mycovirus species as a function of the number of isolates. Yellow box plots represent the species richness values based on linear interpolation of random permutations, and the gray area represents the standard deviation. **(B)** Species richness for mycovirus species in *B. cinerea* isolates collected from raspberry and strawberry plants. **(C)** Shannon and Simpson diversity indexes for mycovirus species (without including the analysis of novel contigs) detected in *B. cinerea* isolates collected from raspberries and strawberries. **(D)** Species co-occurrence matrix to analyze the negative (orange), positive (green), or random (gray) co-occurrences between two mycoviruses and between the mycoviruses and host crop. Mycoviruses with only random associations were excluded from this matrix.

To investigate whether mycovirus species positively or negatively co-occurred with each other or with the *B. cinerea* crop host (strawberry or raspberry), a species co-occurrence matrix was used. Several mycoviruses had positive co-occurrences with other (Fig. 6E); BpBV1 positively co-occurred with Botrytis cinerea fusarivirus 1 (BcFV1), Sclerotinia sclerotiorum deltaflexivirus 2 (SsDFV2), SsDFV2-WX, Monilinia hypovirus A (MHVA), Botrytis cinerea mitovirus 9 (BcMV9), and Botrytis cinerea fusarivirus 3 (BcFV3). BcFV1 also positively co-occurred with Plasmopara viticola lesion associated orfanplasmovirus 5 (PvlaOV5), Grapevine-associated botybirnavirus 1 (GaBV1), and Botrytis virus F (BVF). There were also positive and negative co-occurrences between several mycoviruses and the *B. cinerea* crop host based on the species co-occurrence matrix (Fig. 6E). Botrytis cinerea hypovirus 2 (BcHV2) and Botrytis cinerea fusarivirus 8 (BcFV8) positively co-occurred with isolation from strawberries while negatively co-occurring with isolation from raspberries. Additionally, Botrytis cinerea alpha-like virus 1 (BcAV1) and BpBV1 positively co-occurred with isolation from raspberries while negatively co-occurring with isolation from strawberries. SsDFV2 and the novel virus related to LbBV1 also positively co-occurred with isolation from raspberries.

## Discussion

The use of high-throughput sequencing has allowed for analyses of the diversity of the mycovirome of fungi and for an exponential increase in the number of novel mycoviruses discovered in recent years. Much of the focus of mycovirus research has been on identifying and harnessing mycoviruses as BCA against fungal pathogens. We profiled the mycovirome of 45 isolates of *B. cinerea* collected from fruits in 2019, 2020, and 2021 in Quebec, Canada. To our knowledge, this is the first study to evaluate the mycovirome of a phytopathogenic fungi in Canada. To increase the chances of identifying hypovirulence-inducing mycoviruses, we pre-selected isolates with small or no lesions on detached bean leaves and/or that failed to develop or developed small sclerotia after being grown in the dark on Petri plates for three weeks.

Mycoviruses were found to be highly prevalent, with 44 out of the 45 isolates investigated harboring more than one virus. The species richness was higher than expected, with a total of 81 viral species from 20 classified families and several taxonomic groups unclassified at the family level with dsRNA, ssRNA(+), ssRNA(−), ssDNA, and RT ssRNA genome types. A proportionally higher prevalence of viruses was found than a study evaluating the mycovirome of 248 *B. cinerea* isolates collected from grapevines in Italy and Spain analyzed in 29 pools, which identified 92 mycovirus species. The prevalence of mycoviruses in our study was also greater than studies of the mycovirome of *S. sclerotiorum*; 57 mycoviruses were detected in 85 isolates in Australia, 68 mycoviruses were found in over 300 isolates in China, and 28 mycoviruses were found in 138 isolates in the US These differences could be on account of different analysis methods as these studies used total RNA extraction while we used dsRNA extraction. Additionally, differences in prevalence could be due to pre-selecting isolates with low fitness/pathogenicity rather than randomly selecting isolates. Sampling locations and the crop hosts may also affect the prevalence of mycoviruses. Previous studies have found that there can be an effect of the crop host on mycovirus species richness. For example, a higher percentage of isolates collected from pepper and tomato harbored dsRNA than isolates collected from aubergine and cucumber [38]. In addition, similar percentages of dsRNA elements were reported in *B. cinerea* isolates from grapevine and tomato in Spain as in *B. cinerea* isolates from the same hosts in New Zealand [38, 46].

In the current study, mycovirus species richness and diversity were similar between *B. cinerea* isolates collected from strawberry and raspberry. However, a species co-occurrence matrix suggested that some mycoviruses had a positive co-occurrence with a crop host and had a negative co-occurrence with the other crop host. The species accumulation curve, which did not reach a plateau, indicated that the relatively small sample size limited the capacity of this study to capture the entire diversity of the mycovirus population. As a result, evaluating a larger pool of isolates could reveal more pronounced effects of the crop host. Additionally, most of the *B. cinerea* isolates had low virulence and/or abnormal sclerotial morphotypes which could affect the observed mycovirus diversity. Differences between isolates collected from strawberry and raspberry (two small-fruits plants within *Rosaceae*) might also be less pronounced compared to those collected from more distantly related plant species.

*Endornaviridae*, a family of capsidless viruses with linear ssRNA(+) genomes, was the most prevalent family in this study as over half of the isolates were infected by one or more endornaviruses. These viruses may have contributed to the low fitness/virulence of the isolates as several endornaviruses have been associated with hypovirulence-inducing effects in their hosts [59–61]. One of the putative novel viruses we detected was also a endornavirus, related to Hymenoscyphus fraxineus endornavirus 1. Further, the second most prevalent family was *Botybirnaviridae* which comprises mycoviruses with bipartite dsRNA genomes that can form particles. Only a few botybirnaviruses have been previously identified in *B. cinerea*, including Botrytis cinerea botybirnavirus 1, Botrytis cinerea botybirnavirus 2, and BpBV1. BpBV1– detected in 40% of our isolates–has been associated with hypovirulence-inducing effects. In addition, for the first time in *B. cinerea* isolates, we identified strains of Alternaria botybirnavirus 1, GaBV1, Ipomoea aquatica botybirnavirus, and LbBV1, as well as a putative novel botybirnavirus related to LbBV1.

There have been reports of mycoviruses previously found to be infecting *B. porri*, and *S. sclerotiorum* in *B. cinerea* isolates [30]. In our study, many mycovirus species, including the aforementioned botybirnaviruses, that have been previously detected in different fungal species, such as *Leptosphaeria biglobosa*, *Alternaria alternata*, *Erysiphe necator*, and *Monilinia fructicola*, in addition to *B. porri* and *S. sclerotiorum*, were detected in *B. cinerea*. Given the host range of *B. cinerea* as a generalist species with many host crops, it shares an ecological niche with many other fungal species, therefore making viral cross-species transmission likely [63]. In particular, *B. cinerea* and *S. sclerotiorum* have a high degree of genetic identity and have overlapping host ranges, facilitating the horizontal transfer of mycoviruses [30, 64]. In our *B. cinerea* isolates, we detected 25 species of mycoviruses initially identified in *S. sclerotiorum*. Mycoviruses from other fungal species may be more likely to induce hypovirulence as it has been hypothesized that hypovirulence-inducing mycoviruses have recently infected their hosts [63]. Mycovirus infections are typically asymptomatic as hypovirulence is generally not beneficial to the survival of either the mycovirus or the host fungus [65]. However, mycoviruses that are asymptomatic in one fungus may induce virulence when transmitted to a different fungal species, as they have not yet been able to evolve to cause latent infections [63]. For example, LbBV1 was asymptomatic in *L. biglobosa*, but induced hypovirulence when transmitted to *B. cinerea* [63].

The identification of ssRNA(−) viruses in fungi is rare compared to ssRNA(+) and dsRNA viruses. The first report of a ssRNA(−) mycovirus in *B. cinerea*, Botrytis cinerea negative-stranded RNA virus 1, was in 2016. Subsequent studies, such as those by Hao et al. (2018) and Ruiz-Padilla et al. (2021) [30] revealed the presence of ssRNA(−) mycoviruses in *B. cinerea*, with the latter identifying 15 novel ssRNA(−) species in a survey of 248 isolates. This suggests that ssRNA(−) have been largely undiscovered in *B. cinerea*. In the current study, eight ssRNA(−) mycovirus species were identified, including two putative novel ssRNA(−) species. Further, ssDNA mycoviruses are uncommon in fungi, but BcssDV1, BcHADV1, and BcGV1 were found in two *B. cinerea* isolates in this study. These viruses may contribute to hypovirulence as the ssDNA mycoviruses, BGDaV1, SsHADV-1, and FgGMTV have been reported to have negative effects on the virulence of their hosts, *B. cinerea*, *S. sclerotiorum*, and *F. graminearum*, respectively [15–17]. These mycoviruses are also among the few known mycoviruses to be infectious as viral particles in fungi [15–17]. The capacity of ssDNA to be infectious as particles provides an advantage for their use as BCA as they are not hindered by limitations of vegetative incompatibility.

Although most mycoviruses do not cause any apparent symptoms in their hosts, the majority of the *B. cinerea* isolates selected for analysis in this study had abnormal sclerotial morphotypes and/or low virulence to increase the likelihood of detecting hypovirulence-inducing mycoviruses. The isolate, Bc2019-174, was used to select isolates with small lesion sizes as it produced ‘normal’ lesion sizes. Our analyses revealed that this isolate was not mycovirus-free as a fusarivirus and two partitiviruses were identified. However, it is possible that one or more of these mycoviruses induce hypovirulence as Bc2019-174 failed to produce sclerotia. In addition, several known mycoviruses that induce hypovirulence in *Botrytis* were detected, including BcHV1, BcMV1, and BpBV1 [35, 41, 42]. SsDFV2 and Sclerotinia sclerotiorum narnavirus 4 (SsNV4) identified in this study were also associated with hypovirulence-inducing effects in *S. sclerotiorum*, although their effects on virulence have not been evaluated in *B. cinerea* [66, 67]. While many of the mycoviruses identified might contribute to the reduced virulence and/or abnormal sclerotial morphotype of the *B. cinerea* isolates, the specific effect of any single mycovirus could not be determined within the scope of this study. Since almost every isolate harbored multiple mycoviruses, these mycoviruses could have synergetic, antagonistic, or neutral effects on *B. cinerea*. Future studies will evaluate selected mycoviruses identified in this study for their potential as BCA.

## Conclusions

A high diversity of mycoviruses with ssRNA(+), dsRNA, ssRNA(−), ssDNA, and RT ssRNA genomes were identified in isolates of *B. cinerea*. Hypovirulence-inducing mycoviruses, such as BcMV1, BcHV1, and BpBV1, were identified. In addition, ssDNA viruses in the *Genomoviridae* family that may be infective to *Botrytis* extracellularly were detected. A few mycoviruses positively co-occurred with the isolation of *B. cinerea* from one crop (strawberry or raspberry) and negatively co-occurred with isolation from the other crop. Therefore, there may be an effect of the host crop on the presence of mycoviruses. Many new strains of mycoviruses were identified from *Lenarviricota, Kitrinoviricota, Pisuviricota, Negarnaviricota,* and *Cressdnaviricota* phyla. Furthermore, four putative novel mycovirus species belonging to *Endornaviridae*, *Botybirnaviridae*, *Peribunyaviridae*, and *Bunyavirales* taxa were identified.

Exploratory studies of the regional mycovirus populations are important for the targeted use of mycoviruses as BCA and to minimize the risk of introducing the mycoviruses BCA as harmful exotic organisms. Moving forward, mycoviruses that produce viral particles that can be applied exogenously appear to hold significant promise as durable BCA against *B. cinerea*. To enhance our understanding of the novel mycoviruses with potential as BCA identified in this study, it will be essential to employ diverse methodologies, such as cycloheximide and ribavirin treatments, hyphal tip isolation, protoplast regeneration, and single-spore hybridization [68]. These approaches coupled with transcriptomic and phenotypic analyses could help evaluate the restoration of normal phenotypes, such as sclerotia production, and further elucidate the interactions between mycoviruses and their hosts. Ultimately, a comprehensive assessment of these mycoviruses may pave the way for innovative biocontrol strategies that effectively mitigate *B. cinerea* in various agricultural settings.

## Materials and Methods

### Botrytis cinerea isolate collection

A total of 129 isolates of *B. cinerea* were collected from strawberry, raspberry, and grapevine plants from seven commercial and experimental farms in the Montérégie and Estrie regions in Quebec, Canada in 2019, 2020, and 2021. One sample was taken per plant with a sampling swab and the swab was rubbed onto a PDA plate which was kept at room temperature. Once the colony started to develop, the isolates were subcultured onto new PDA plates. After the colony had grown (usually after 4–5 days), one agar plug per isolate was placed in sterilized soil in 2-mL tubes. Once the isolates colonized the soil, the tubes were stored at 2–4°C until further processing. The isolates collected in 2019 were later subcultured onto PDA plates and stored at 4°C until further processing.

### Botrytis cinerea isolate morphotype/virulence evaluation

Isolates were evaluated based on virulence as represented by lesion size on bean (*Phaseolus vulgaris* L.) leaves [69] and were placed into colony morphotype categories as in Jia et al. (2021) and Martinez et al. (2003) [69]. To evaluate virulence, the leaves were sterilized in a 0.5% bleach solution for 30 sec, rinsed twice in distilled water, and dried with autoclaved cotton paper. The isolates were grown on 9-cm PDA plates for four days. Then, 5-mm agar plugs were transferred to the sterilized bean leaves. The leaves were placed on PDA plates in a growth cabinet with a 12-h photoperiod at 20°C for 72 h. Lesion size was measured with a caliper. Three to six replicates per isolate were evaluated. To determine the colony morphotype, the isolates were grown on 9-cm PDA plates, and 5-mm agar plugs were transferred to new PDA plates, covered with aluminum foil, and grown in the dark for three weeks. Isolates were then classified into sclerotial type and/or mycelial type categories according to Martinez et al. (2003) [69] (Fig. 1A). Sclerotial types were classified as follows: S1, sclerotia on the edge of the Petri dish; S2, predominately large sclerotia in a circle; S3, predominately large sclerotia with irregular placements; and S4, numerous small sclerotia with irregular placements. Mycelial types were classified as follows: M1, short mycelium without conidiation; M2, aerial mycelium with conidiation; M3, mycelial masses with or without conidiation; and M4, thick and woolly mycelium without conidition [69].

Of the 45 selected isolates, seven isolates were collected from raspberries in Oka, 15 were from raspberries in Shefford, 19 were from strawberries in Sainte-Anne-des-Plaines, one was from a strawberry plant in Sainte-Catherine-de-Hatley, one was from a strawberry plant in Saint-Malo, one was from a strawberry plant in Saint-Paul-d’Abbotsford, and one was from grapevine from Saint-Bernard-de-Lacolle (Supplementary Table 1; Fig. 7).

**Fig 7.**
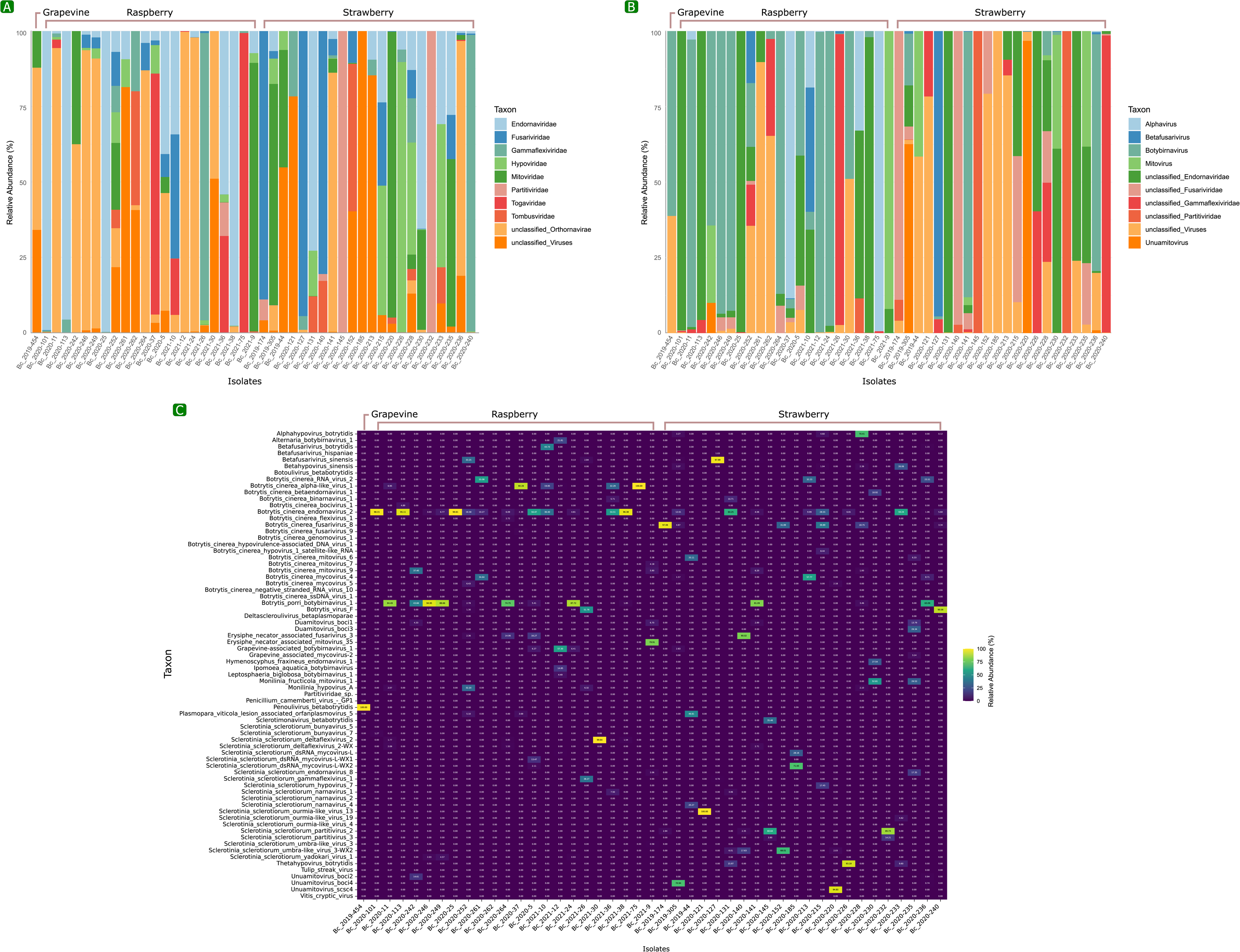
Collection sites in Quebec, Canada of *Botrytis cinerea* isolates. The color corresponds to the year in which the samples were collected, and the size of the bubble corresponds to the number of isolates collected from a site.

### B. cinerea sample preparation for dsRNA extraction

The isolates selected for sequencing were grown on 9-cm PDA plates. After seven days, agar plugs were transferred to new PDA plates covered with sterile cellophane and maintained for at least seven days or until sufficient mycelium had grown. Mycelia were harvested and stored at - 80°C. The samples were placed in 25-mL mixer grinding jars containing a single 20-mm stainless steel grinding ball pre-chilled with liquid nitrogen and homogenized into a powder with a Mixer Mill MM 400 (Retsch, Haan, Germany) at a frequency of 30 Hz for 30 sec. The samples were returned to −80°C.

### dsRNA extraction

In this study, we explored various published dsRNA extraction methods for detecting and characterizing mycoviruses. However, despite multiple attempts, the quantity of dsRNA obtained was consistently low, and the results were inconsistent across different trials. To address this issue, refined a dsRNA extraction method for mycovirus detection and characterization, leveraging our expertise in RNA and dsRNA extraction [52, 70–73]. In particular, our methods were based on a cellulose column chromatography method in Poursalavati et al. (2023).

First, 2.4 mL of TRIzol Reagent (Invitrogen, Carlsbad, CA) and 30 µL of mercaptoethanol (Millipore-Sigma, Oakville, Canada) were added to 5-mL macrotubes containing 0.6 g of mycelium. Phaseolus vulgaris endornavirus 1 (PvEV-1), a virus of common bean, was used as a positive control for dsRNA. To produce the positive control, bean leaves infected with PvEV-1 were homogenized into a powder using liquid nitrogen and kept at −80°C. The infected leaves were suspended in 800 µL TRIzol and centrifuged at 12,298 × *g* at 4°C for 10 min. The supernatant was transferred to a new tube and added to each 5-mL tube to constitute 1% of the sample. Next, 900 µL of chloroform:isoamyl alcohol 24:1 (Millipore-Sigma) was added. The tubes were vortexed and centrifuged for 10 min at 12,298 × *g* at 4°C. The supernatant was transferred into new 5-mL macrotubes. An equal volume of chloroform:isoamyl alcohol 24:1 was added, and the tubes were centrifuged for 10 min at 12,298 × *g* at 4°C. The supernatant was transferred into new 5-mL tubes.

Second, cellulose column chromatography was performed to isolate the dsRNA. A cellulose slurry with 200 mg of cellulose (Sigmacell Cellulose, Type 101, Millipore-Sigma) per mL of a 16-S chromatography buffer [50 mM Tris; 100 mM NaCl, pH 6.8, containing 16% (*v/v*) ethanol] was vortexed and gently mixed on an orbital shaker at 2,000 rpm until the cellulose was homogenized. Ethanol was added to each sample to reach 16% ethanol. A 1/10 volume of chromatography buffer (500 mM Tris, 1 M NaCl, pH 6.8) and 500 µL of cellulose slurry were also added. The samples were vortexed and then shaken on an orbital shaker at 2,000 rpm for 15 min. The tubes were centrifuged at 17,709 × *g* for 1 min and the supernatant was discarded. A 1.5 mL volume of the 16-S chromatography buffer was added, vortexed, and shaken on an orbital shaker for 10 min. The tubes were centrifuged at 17,709 × *g* for 1 min. The supernatant was discarded and 500 µL of the 16-S chromatography buffer was added and mixed by pipetting. A 500 µL volume of each sample was added to 0.22-µm filter mini columns (Norgen Biotek Corporation, Thorold, Canada), which were centrifuged at 10,000 × *g* for 10 sec. A volume of 500 µL of each sample was added to the columns and mixed by pipetting. The columns were centrifuged at 10,000 × *g* and the flow-through was discarded. This step was repeated until all the sample was processed. Next, 500 µL of the 16-S chromatography buffer was added to each column and centrifuged at 10,000 × *g* for 10 sec. This step was repeated. Then, the columns were dried by centrifuging at 10,000 × *g* for 10 sec. The columns were placed in new tubes, 400 µL of an elution buffer (50 mM Tris; 100 mM NaCl, pH 6.8) was added and the tubes were centrifuged at 10,000 × *g* for 10 sec. Next, 4 M of LiCl and one volume of isopropyl palmitate (Bioshop) were added. The tubes were kept at −20°C for 2 h.

The dsRNAs were precipitated at 25,155 × *g* for 20 min and washed twice with 70% ethanol. The pellet was suspended in 30 µL of ultrapure water. Then, nucleic acids were quantified with a Qubit 4.0 fluorometer with Qubit dsDNA HS Assay Kit and Qubit RNA HS Assay Kit (Invitrogen). The samples were digested with DNase 1 (Thermo Fisher Scientific Nepean, Canada or New England BioLabs, Whitby, Canada) and RNase T1 (Thermo Fisher Scientific) based on the concentration of dsDNA and RNA in the presence of a 10x React Buffer with MgCl_2_ (Thermo Fisher Scientific).

Ultrapure water was added to reach 200 µL and the enzymes were inactivated by adding an equal volume of phenol:chloroform:isoamyl alcohol (Thermo Fisher Scientific), vortexing, and centrifuging for 10 min at 12,298 × *g* at 4°C. The supernatant was transferred to new tubes and an equal volume of chloroform:isoamyl alcohol 24:1 was added and the tubes were centrifuged for 10 min at 25,155 × *g* at 4°C. Two volumes of 100% ethanol and 1/10 volume of sodium acetate (3M, pH 5.2) were added and the tubes were kept at −20°C overnight. The tubes were precipitated at 25,155 × *g* for 20 min, washed twice with 80% ethanol, and resuspended in 20 µL of ultrapure water. The dsRNAs were quantified with a Qubit 4.0 fluorometer with Qubit dsDNA HS Assay Kit and a Qubit RNA HS Assay Kit to ensure that the RNA was fully digested.

### dscDNA synthesis and sequencing

Viral double-stranded cDNA (dscDNA) synthesis was conducted according to Fall et al. (2020) [70] by mixing 5 μL of dsRNA with 60 μM of random primers (NEB), 10 nM of dNTP (NEB), and 2.5 μL of nuclease-free water. The mixture was heated at 99°C for 5 min and then immediately placed on ice. To synthesize the first strand of dscDNA, 4 μL of first strand buffer, 1 μL of 0.1 M DTT, 0.5 μL of RNase inhibitor, murine (40U/μl) (NEB), and 1 μL of Superscript III (Invitrogen) were added. The mixture was incubated at 25°C for 5 min for priming, then at 50°C for 45 min. The reaction was inactivated by heating at 70°C for 15 min. Next, 1 μL of RNase H (NEB) was added for the removal of RNA in the DNA-RNA hybrid. The mixture was incubated at 37°C for 20 min. To synthesize the second strand of dscDNA, 2.5 μL of klenow 10x buffer supplemented with NAD, 1 μL of 10 nM of dNTP, 0.5 μL of Klenow DNA polymerase I (2.5 U) (Promega, Madison, USA), and 1 μL of *E. coli* DNA ligase 1 (10 U) (NEB) were added. The reaction was incubated at 16°C for 150 min, and the ligase was inactivated at 65°C for 20 min.

The samples were cleaned with Agencourt AMPure XP magnetic beads (Beckman-Coulter), and the dscDNA was quantitated using a Qubit 4 fluorometer (Thermo Fisher Scientific) and adjusted to a concentration of 0.2 ng/µl. One ng of dscDNA was used for library preparation with the Nextera XT DNA Library Preparation Kit (Illumina, San Diego, USA), following the manufacturer’s instructions. The quality of the libraries was verified using the High Sensitivity DNA Reagents Kit with the Agilent 2100 Bioanalyzer system (Santa, Clara, USA). The prepared libraries were pooled in an equimolar ratio to a final concentration of 4 nM. The denatured pooled libraries were then diluted to 12 pM before loading onto the reagent cartridge of the MiSeq Reagent Kit v3. Sequencing was performed with 2x 301 cycles using a paired-end read type on the MiSeq System (Illumina).

### Bioinformatic analyses

Bioinformatics analyses were carried out using an in-house pipeline called SOVAP [52, 74]. Briefly, Fastp v.023.2 [75] was used to remove low-quality reads and Illumina sequence adapters from raw sequencing data, and Centrifuge v1.04 [76] was utilized to decontaminate the reads in tandem with a custom-built database [77], effectively eliminating traces of bacterial, archaeal, and fungal genomes (--min-hitlen 50). SPAdes v3.15.5 [78] was used to assemble the contigs (-- rna), and geNomad v.1.3.3 [79] was used to identify the viral contigs (end-to-end -s 7.0). Contigs of > 500 bp in length were clustered with CD-HIT v.4.8.1 (-c 0.95 -n 10 -aS 0.85) [80, 81], and the relative abundance of viral contigs was estimated for each contig using BWA v.0.7.17-r1188 [82], followed by calculation of relative abundance values through an in-house script [83]. DIAMOND (blastx) v.2.1.4 (--sensitive) [84] was employed for annotating clustered viral contigs, utilizing custom-built GenBank viral genomes (accessed on 2023.11.07) alongside an in-house indexed IMG/VR v4 database [85].

For phylogenetic analyses to identify novel viruses, the RNA-dependent RNA polymerase (RdRp) gene was used as a hallmark gene to identify RNA viruses, while the rolling-circle replication initiation protein (Rep) was used to identify ssDNA viruses [86]. RdRp genes were identified from the viral proteins (> 70 amino acids) reported in geNomad with Diamond Blastp (--very-sensitive) and RdRp-Scan datasets [52, 87]. The RdRp-Scan [87] and NeoRdRp [88] Hidden Markov model (HMM) profile database using HMMSearch v3.3.2 was used to re-analyze the resulting hits. Contigs with E values <0.001 and scores >50 were clustered with CD-HIT to discard contigs with 95% average amino acid identity. The longest contig candidates from each cluster were classified into taxonomic ranks for *Lenarviricota*, *Kitrinoviricota*, *Pisuviricota*, *Negarnaviricota*, *Botybirnaviridae*, and taxonomic groups that are unclassified at the phylum level. The candidates underwent alignment via Clustal Omega v1.2.4 (--auto) against the pre-built backbone trees of RdRp-Scan, followed by tree generation utilizing FastTreeMP v2.1.11 with multi-threading (-wag -spr 4 -mlacc 2 -pseudo -slownni). Subsequently, the phylogenetic trees were rendered and visualized using iToL v6.7.3.

Then, a multi-step methodology was employed to re-examine RdRp candidates and uncover novel sequences. Initially, the three key RdRp motifs (A, B, and C) were identified using CLC Genomics Workbench v22 (QIAGEN, Hilden, Germany) for the *Lenarviricota*, *Kitrinoviricota*, *Pisuviricota*, and *Negarnaviricota* phyla and within the family *Botybirnaviridae*. Complementary literature searches were conducted to determine motifs for DNA viruses and RNA viruses that are unclassified at the phylum level. Candidates possessing all the motifs were selected, followed by multiple-sequence alignments (MSAs) analysis utilizing MAFFT via MegAlign Pro v17.5.0 (DNAStar Lasergene software) to discern conserved regions and potential variations among the candidates within each taxon. A manual curation step ensued post-MSA, wherein candidates with non-perfectly aligned motifs were eliminated. Subsequently, the phylogenetic trees were reconstructed, retaining only those candidates containing all motifs as previously described. Finally, RdRp and Rep sequences of the confirmed candidates were submitted to GenBank for validation and the associated sequencing raw data were submitted into the NCBI sequence read archive database (SRA).

### Statistical analyses

A map of sampling locations, an accumulation curve, a co-occurrence analysis, Shannon diversity index analysis, and a plot of the number of occurrences of mycovirus species in *B. cinerea* isolates were created in R version 4.2.3 with the following packages: ggplot2, ggmap, tidyverse, googlesheets4, ggpubr, lubridate, openintro, maps, ggmap, ggthemes, multcomp, cooccur, reshape2, dplyr, tidyr, and vegan.

The analysis of radial growth of *B. cinerea* isolates compared to a control isolate, Bc2019-174, was conducted using Dunnett’s test. A Type 1 error of *P* = 0.05 was set for all statistical tests. To characterize the mycovirome of the isolates, including relative abundance of taxon, species richness, and Shannon and Simpson diversity indexes, the positive control (PvEV-1), phages, and other obvious non-mycovirus taxa (Supplementary Table 1) were excluded. Shannon diversity and Simpson diversity indexes were calculated using the transcripts per million (TPM) values of each mycovirus species. These diversity indexes excluded the analysis of novel contigs. A t-test was performed to test for differences in species richness and in the Shannon and Simpson diversity indexes between *B. cinerea* isolates from raspberries and strawberries. A species co-occurrence analysis was used to evaluate the negative, positive, and random co-occurrences between mycoviruses, and between the mycoviruses and the host crop.

### Data availability statement

All the raw sequencing reads are available at the NCBI SRA database: BioProject accession no. PRJNA1096650, BioSample accession numbers SAMN40758340 to SAMN40758384, and SRA accession numbers SRR28563938 to SRR28563899 (https://dataview.ncbi.nlm.nih.gov/object/PRJNA1096650?reviewer=d94em0mmn1f9lv3sjjvihcs v5v). The Genbank accession numbers are in Table 1.

## Acknowledgements

The authors gratefully acknowledge Annie Lefebvre and Pierre-Olivier Hebert, research assistants in the fungal epidemiology lab at the Saint-Jean-sur-Richelieu Research and Development Centre, Agriculture and Agri-Food Canada, for their help and assistance. We are also grateful to Kamal Bouarab and Isabelle Laforest-Lapointe from Université de Sherbrooke for their support and advice. We would also like to acknowledge the administrative support that we received from the team at the Saint-Jean-sur-Richelieu Research and Development Centre (the teams led by Vicky Toussaint and Mélanie Maheux in particular).

