## Supplementary materials for "Exploring the mycovirome: novel and diverse mycoviruses in *Botrytis cinerea*"

**Supplementary material**

**Table 1** Collection information, colony morphotype and lesion size of the 45 selected *Botrytis cinerea* isolates from Quebec, Canada (2019, 2020, and 2021).

| Isolate id | Host | Site | Year | Colony morphotype[1] | Lesion size ($mm^2$) |
|---|---|---|---|---|---|
| **Bc2019-44** | Strawberry | Sainte-Catherine-de-Hatley | 2019 | S4 | 424 |
| **Bc2019-174** | Strawberry | Saint-Malo | 2019 | M2 | 756 |
| **Bc2019-305** | Strawberry | Saint-Paul-d'Abbotsford | 2019 | M2/S4 | 169 |
| **Bc2019-454** | Grapevine | Saint-Bernard-de-Lacolle | 2019 | M1 | 81 |
| **Bc2020-5** | Raspberry | Shefford | 2020 | S3 | 1005 |
| **Bc2020-11** | Raspberry | Shefford | 2020 | S3 | 29 |
| **Bc2020-25** | Raspberry | Shefford | 2020 | S4 | 46 |
| **Bc2020-37** | Raspberry | Shefford | 2020 | S3 | 0 |
| **Bc2020-101** | Raspberry | Shefford | 2020 | M3/S3 | 185 |
| **Bc2020-113** | Raspberry | Shefford | 2020 | S3 | 14 |
| **Bc2020-121** | Strawberry | Sainte-Anne-des-Plaines | 2020 | S3 | 265 |
| **Bc2020-127** | Strawberry | Sainte-Anne-des-Plaines | 2020 | M1 | 183 |
| **Bc2020-131** | Strawberry | Sainte-Anne-des-Plaines | 2020 | S2 | 165 |
| **Bc2020-140** | Strawberry | Sainte-Anne-des-Plaines | 2020 | S3 | 242 |
| **Bc2020-141** | Strawberry | Sainte-Anne-des-Plaines | 2020 | S4 | 74 |
| **Bc2020-145** | Strawberry | Sainte-Anne-des-Plaines | 2020 | M3/S3 | 301 |
| **Bc2020-152** | Strawberry | Sainte-Anne-des-Plaines | 2020 | S2 | 126 |
| **Bc2020-185** | Strawberry | Sainte-Anne-des-Plaines | 2020 | M3 | 15 |
| **Bc2020-213** | Strawberry | Sainte-Anne-des-Plaines | 2020 | S3 | 118 |
| **Bc2020-215** | Strawberry | Sainte-Anne-des-Plaines | 2020 | S4 | 108 |
| **Bc2020-220** | Strawberry | Sainte-Anne-des-Plaines | 2020 | S3 | 9 |
| **Bc2020-226** | Strawberry | Sainte-Anne-des-Plaines | 2020 | S4 | 15 |

| | | | | | |
|---|---|---|---|---|---|
| **Bc2020-228** | Strawberry | Sainte-Anne-des-Plaines | 2020 | S3 | 180 |
| **Bc2020-230** | Strawberry | Sainte-Anne-des-Plaines | 2020 | S2 | 276 |
| **Bc2020-232** | Strawberry | Sainte-Anne-des-Plaines | 2020 | S3 | 287 |
| **Bc2020-233** | Strawberry | Sainte-Anne-des-Plaines | 2020 | S4 | 29 |
| **Bc2020-235** | Strawberry | Sainte-Anne-des-Plaines | 2020 | S3 | 264 |
| **Bc2020-236** | Strawberry | Sainte-Anne-des-Plaines | 2020 | -[3] | 0 |
| **Bc2020-240** | Strawberry | Sainte-Anne-des-Plaines | 2020 | S4 | 211 |
| **Bc2020-242** | Raspberry | Oka | 2020 | S3 | 18 |
| **Bc2020-246** | Raspberry | Oka | 2020 | S2 | 199 |
| **Bc2020-249** | Raspberry | Oka | 2020 | S3 | 0 |
| **Bc2020-252** | Raspberry | Oka | 2020 | S3 | 14 |
| **Bc2020-261** | Raspberry | Oka | 2020 | S2 | 279 |
| **Bc2020-262** | Raspberry | Oka | 2020 | S2 | 113 |
| **Bc2020-264** | Raspberry | Oka | 2020 | S3 | 307 |
| **Bc2021-9** | Raspberry | Shefford | 2021 | S2 | 706 |
| **Bc2021-10** | Raspberry | Shefford | 2021 | S4 | 308 |
| **Bc2021-12** | Raspberry | Shefford | 2021 | S1 | 61 |
| **Bc2021-24** | Raspberry | Shefford | 2021 | S3 | 322 |
| **Bc2021-26** | Raspberry | Shefford | 2021 | S3 | 14 |
| **Bc2021-30** | Raspberry | Shefford | 2021 | S3 | 200 |
| **Bc2021-36** | Raspberry | Shefford | 2021 | M1 | 415 |
| **Bc2021-38** | Raspberry | Shefford | 2021 | S3 | 440 |
| **Bc2021-75** | Raspberry | Shefford | 2021 | M1 | 0 |

[1]Colony morphotypes after growing isolates on potato dextrose agar Petri plates in the dark for three weeks. Classifications were based on four mycelial types and/or four sclerotial types as in Martinez et al. (2003) [69].
[2]Lesion size of *B. cinerea* isolates was measured on detached bean leaves.
[3]Isolate did not grow so did not produce a morphotype.

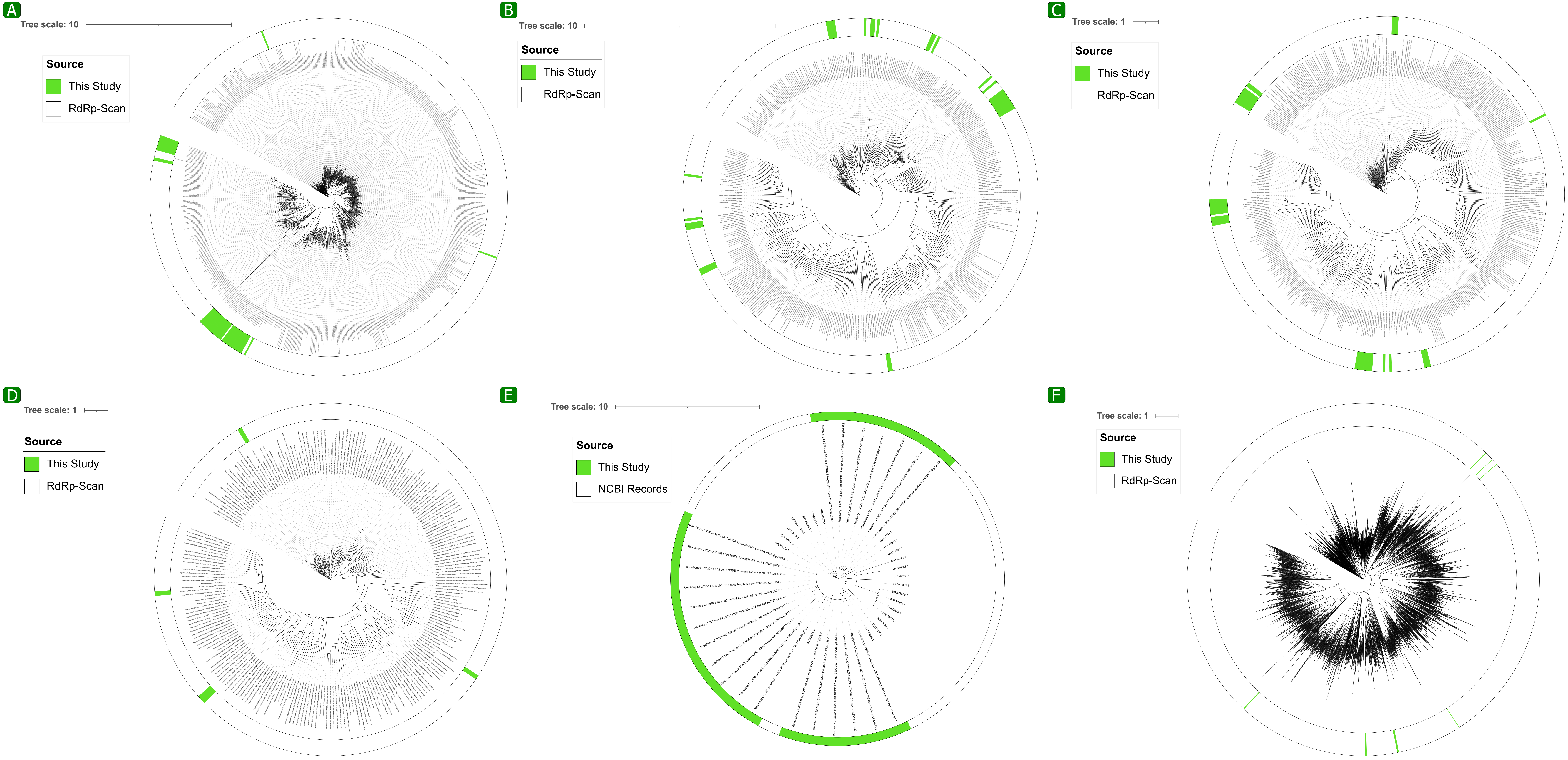
A
Tree scale: 10
Source
This Study
RdRp-Scan
B
Tree scale: 10
Source
This Study
RdRp-Scan
C
Tree scale: 1
Source
This Study
RdRp-Scan
D
Tree scale: 1
Source
This Study
RdRp-Scan
E
Tree scale: 10
Source
This Study
NCBI Records
F
Tree scale: 1
Source
This Study
RdRp-Scan

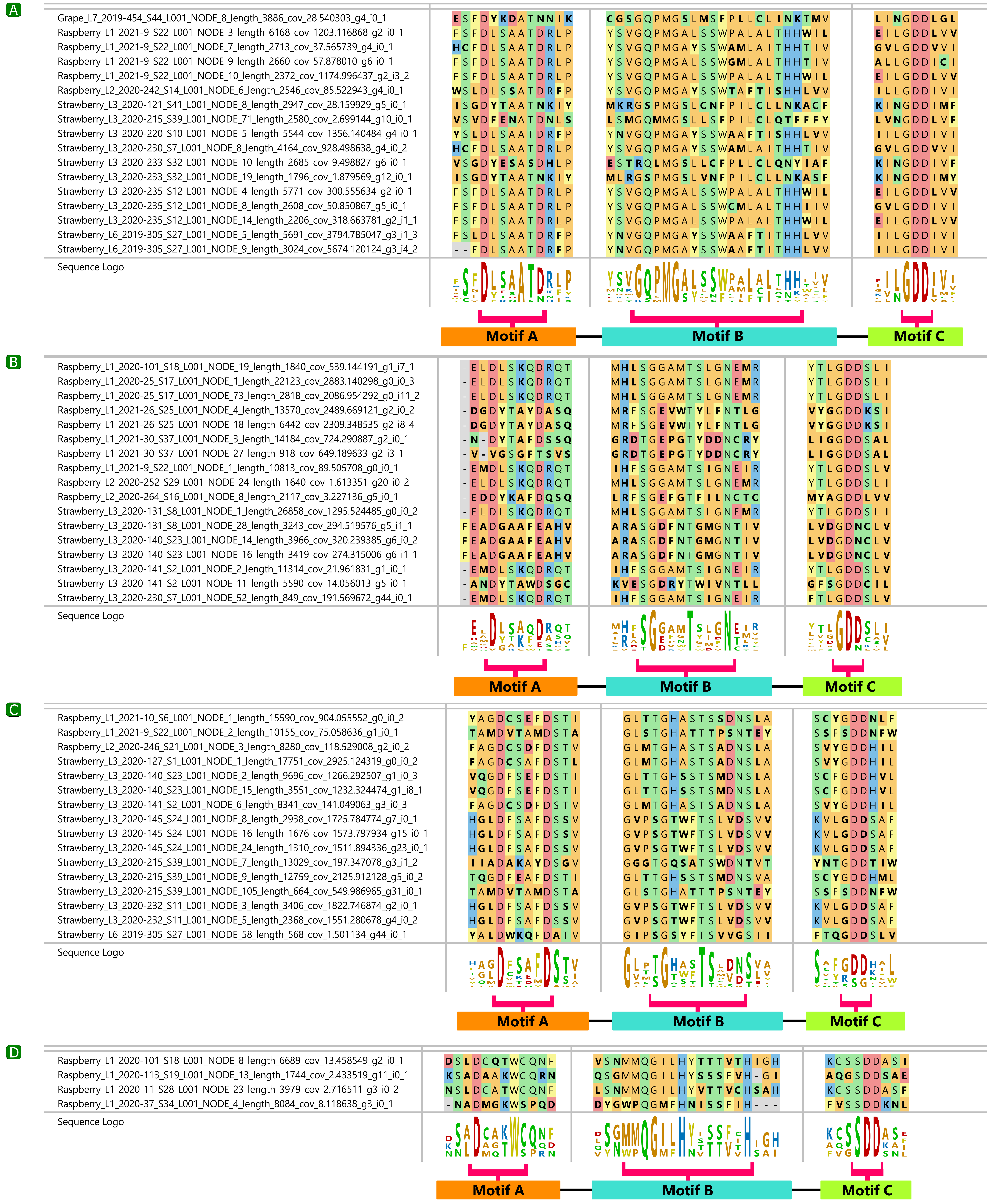
A
Grape_L7_2019-454_S44_L001_NODE_8_length_3886_cov_28.540303_g4_i0_1
Raspberry_L1_2021-9_S22_L001_NODE_3_length_6168_cov_1203.116868_g2_i0_1
Raspberry_L1_2021-9_S22_L001_NODE_7_length_2713_cov_37.565739_g4_i0_1
Raspberry_L1_2021-9_S22_L001_NODE_9_length_2660_cov_57.878010_g6_i0_1
Raspberry_L1_2021-9_S22_L001_NODE_10_length_2372_cov_1174.996437_g2_i3_2
Raspberry_L2_2020-242_S14_L001_NODE_6_length_2546_cov_85.522943_g4_i0_1
Strawberry_L3_2020-121_S41_L001_NODE_8_length_2947_cov_28.159929_g5_i0_1
Strawberry_L3_2020-215_S39_L001_NODE_71_length_2580_cov_2.699144_g10_i0_1
Strawberry_L3_2020-220_S10_L001_NODE_5_length_5544_cov_1356.140484_g4_i0_1
Strawberry_L3_2020-230_S7_L001_NODE_8_length_4164_cov_928.498638_g4_i0_2
Strawberry_L3_2020-233_S32_L001_NODE_10_length_2685_cov_9.498827_g6_i0_1
Strawberry_L3_2020-233_S32_L001_NODE_19_length_1796_cov_1.879569_g12_i0_1
Strawberry_L3_2020-235_S12_L001_NODE_4_length_5771_cov_300.555634_g2_i0_1
Strawberry_L3_2020-235_S12_L001_NODE_8_length_2608_cov_50.850867_g5_i0_1
Strawberry_L3_2020-235_S12_L001_NODE_14_length_2206_cov_318.663781_g2_i1_1
Strawberry_L6_2019-305_S27_L001_NODE_5_length_5691_cov_3794.785047_g3_i1_3
Strawberry_L6_2019-305_S27_L001_NODE_9_length_3024_cov_5674.120124_g3_i4_2
Sequence Logo
Motif A
Motif B
Motif C
B
Raspberry_L1_2020-101_S18_L001_NODE_19_length_1840_cov_539.144191_g1_i7_1
Raspberry_L1_2020-25_S17_L001_NODE_1_length_22123_cov_2883.140298_g0_i0_3
Raspberry_L1_2020-25_S17_L001_NODE_73_length_2818_cov_2086.954292_g0_i11_2
Raspberry_L1_2021-26_S25_L001_NODE_4_length_13570_cov_2489.669121_g2_i0_2
Raspberry_L1_2021-26_S25_L001_NODE_18_length_6442_cov_2309.348535_g2_i8_4
Raspberry_L1_2021-30_S37_L001_NODE_3_length_14184_cov_724.290887_g2_i0_1
Raspberry_L1_2021-30_S37_L001_NODE_27_length_918_cov_649.189633_g2_i3_1
Raspberry_L1_2021-9_S22_L001_NODE_1_length_10813_cov_89.505708_g0_i0_1
Raspberry_L2_2020-252_S29_L001_NODE_24_length_1640_cov_1.613351_g20_i0_2
Raspberry_L2_2020-264_S16_L001_NODE_8_length_2117_cov_3.227136_g5_i0_1
Strawberry_L3_2020-131_S8_L001_NODE_1_length_26858_cov_1295.524485_g0_i0_2
Strawberry_L3_2020-131_S8_L001_NODE_28_length_3243_cov_294.519576_g5_i1_1
Strawberry_L3_2020-140_S23_L001_NODE_14_length_3966_cov_320.239385_g6_i0_2
Strawberry_L3_2020-140_S23_L001_NODE_16_length_3419_cov_274.315006_g6_i1_1
Strawberry_L3_2020-141_S2_L001_NODE_2_length_11314_cov_21.961831_g1_i0_1
Strawberry_L3_2020-141_S2_L001_NODE_11_length_5590_cov_14.056013_g5_i0_1
Strawberry_L3_2020-230_S7_L001_NODE_52_length_849_cov_191.569672_g44_i0_1
Sequence Logo
Motif A
Motif B
Motif C
C
Raspberry_L1_2021-10_S6_L001_NODE_1_length_15590_cov_904.055552_g0_i0_2
Raspberry_L1_2021-9_S22_L001_NODE_2_length_10155_cov_75.058636_g1_i0_1
Raspberry_L2_2020-246_S21_L001_NODE_3_length_8280_cov_118.529008_g2_i0_2
Strawberry_L3_2020-127_S1_L001_NODE_1_length_17751_cov_2925.124319_g0_i0_2
Strawberry_L3_2020-140_S23_L001_NODE_2_length_9696_cov_1266.292507_g1_i0_3
Strawberry_L3_2020-140_S23_L001_NODE_15_length_3551_cov_1232.324474_g1_i8_1
Strawberry_L3_2020-141_S2_L001_NODE_6_length_8341_cov_141.049063_g3_i0_3
Strawberry_L3_2020-145_S24_L001_NODE_8_length_2938_cov_1725.784774_g7_i0_1
Strawberry_L3_2020-145_S24_L001_NODE_16_length_1676_cov_1573.797934_g15_i0_1
Strawberry_L3_2020-145_S24_L001_NODE_24_length_1310_cov_1511.894336_g23_i0_1
Strawberry_L3_2020-215_S39_L001_NODE_7_length_13029_cov_197.347078_g3_i1_2
Strawberry_L3_2020-215_S39_L001_NODE_9_length_12759_cov_2125.912128_g5_i0_2
Strawberry_L3_2020-215_S39_L001_NODE_105_length_664_cov_549.986965_g31_i0_1
Strawberry_L3_2020-232_S11_L001_NODE_3_length_3406_cov_1822.746874_g2_i0_1
Strawberry_L3_2020-232_S11_L001_NODE_5_length_2368_cov_1551.280678_g4_i0_2
Strawberry_L6_2019-305_S27_L001_NODE_58_length_568_cov_1.501134_g44_i0_1
Sequence Logo
Motif A
Motif B
Motif C
D
Raspberry_L1_2020-101_S18_L001_NODE_8_length_6689_cov_13.458549_g2_i0_1
Raspberry_L1_2020-113_S19_L001_NODE_13_length_1744_cov_2.433519_g11_i0_1
Raspberry_L1_2020-11_S28_L001_NODE_23_length_3979_cov_2.716511_g3_i0_2
Raspberry_L1_2020-37_S34_L001_NODE_4_length_8084_cov_8.118638_g3_i0_1
Sequence Logo
Motif A
Motif B
Motif C

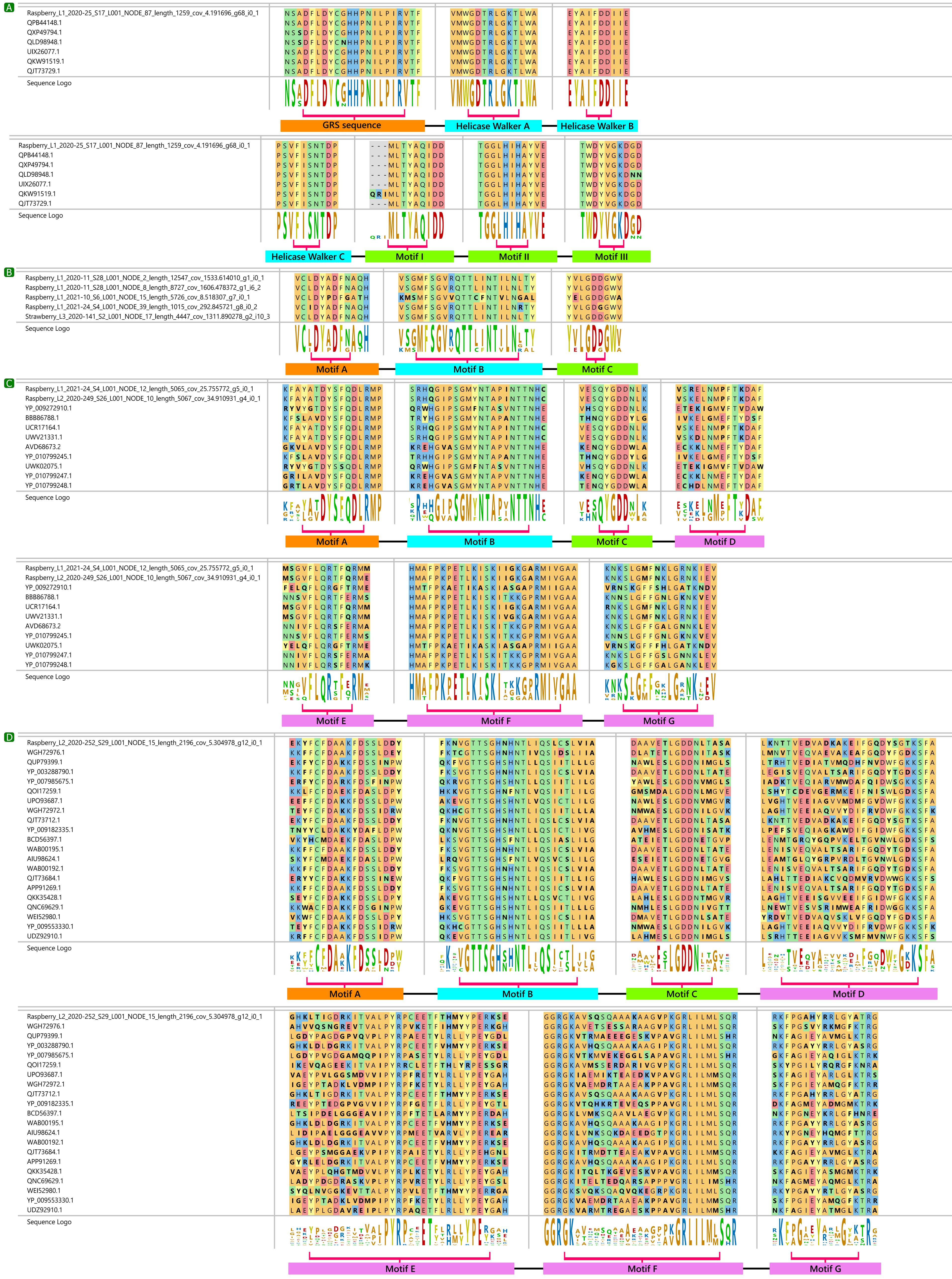
A
Raspberry_L1_2020-25_S17_L001_NODE_87_length_1259_cov_4.191696_g68_i0_1
QPB44148.1
QXP49794.1
QLD98948.1
UIX26077.1
QKW91519.1
QJT73729.1
Sequence Logo
GRS sequence
Helicase Walker A
Helicase Walker B
Helicase Walker C
Motif I
Motif II
Motif III
B
Raspberry_L1_2020-11_S28_L001_NODE_2_length_12547_cov_1533.614010_g1_i0_1
Raspberry_L1_2020-11_S28_L001_NODE_8_length_8727_cov_1606.478372_g1_i6_2
Raspberry_L1_2021-10_S6_L001_NODE_15_length_5726_cov_8.518307_g7_i0_1
Raspberry_L1_2021-24_S4_L001_NODE_39_length_1015_cov_292.845721_g8_i0_2
Strawberry_L3_2020-141_S2_L001_NODE_17_length_4447_cov_1311.890278_g2_i10_3
Motif A
Motif B
Motif C
C
Raspberry_L1_2021-24_S4_L001_NODE_12_length_5065_cov_25.755772_g5_i0_1
Raspberry_L2_2020-249_S26_L001_NODE_10_length_5067_cov_34.910931_g4_i0_1
YP_009272910.1
BBB86788.1
UCR17164.1
UWV21331.1
AVD68673.2
YP_010799245.1
UWK02075.1
YP_010799247.1
YP_010799248.1
Motif D
Motif E
Motif F
Motif G
D
Raspberry_L2_2020-252_S29_L001_NODE_15_length_2196_cov_5.304978_g12_i0_1
WGH72976.1
QUP79399.1
YP_003288790.1
YP_007985675.1
QOI17259.1
UPO93687.1
WGH72972.1
QJT73712.1
YP_009182335.1
BCD56397.1
WAB00195.1
AIU98624.1
WAB00192.1
QJT73684.1
APP91269.1
QKK35428.1
QNC69629.1
WEI52980.1
YP_009553330.1
UDZ92910.1

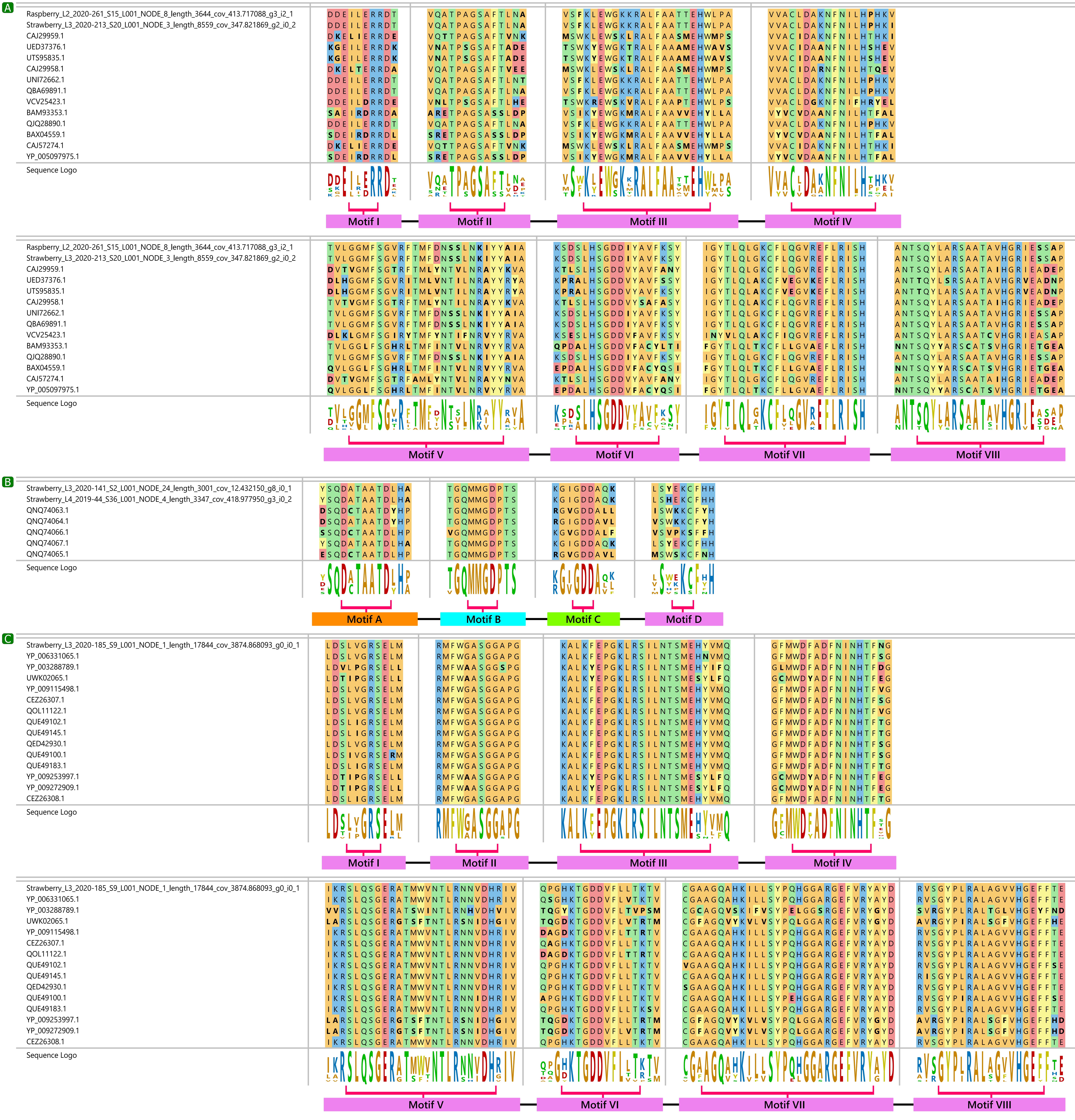
A
Raspberry_L2_2020-261_S15_L001_NODE_8_length_3644_cov_413.717088_g3_i2_1
Strawberry_L3_2020-213_S20_L001_NODE_3_length_8559_cov_347.821869_g2_i0_2
CAJ29959.1
UED37376.1
UTS95835.1
CAJ29958.1
UNI72662.1
QBA69891.1
VCV25423.1
BAM93353.1
QJQ28890.1
BAX04559.1
CAJ57274.1
YP_005097975.1
Sequence Logo
Motif I
Motif II
Motif III
Motif IV
Motif V
Motif VI
Motif VII
Motif VIII
B
Strawberry_L3_2020-141_S2_L001_NODE_24_length_3001_cov_12.432150_g8_i0_1
Strawberry_L4_2019-44_S36_L001_NODE_4_length_3347_cov_418.977950_g3_i0_2
QNQ74063.1
QNQ74064.1
QNQ74066.1
QNQ74067.1
QNQ74065.1
Sequence Logo
Motif A
Motif B
Motif C
Motif D
C
Strawberry_L3_2020-185_S9_L001_NODE_1_length_17844_cov_3874.868093_g0_i0_1
YP_006331065.1
YP_003288789.1
UWK02065.1
YP_009115498.1
CEZ26307.1
QOL11122.1
QUE49102.1
QUE49145.1
QED42930.1
QUE49100.1
QUE49183.1
YP_009253997.1
YP_009272909.1
CEZ26308.1
Sequence Logo
Motif I
Motif II
Motif III
Motif IV
Motif V
Motif VI
Motif VII
Motif VIII

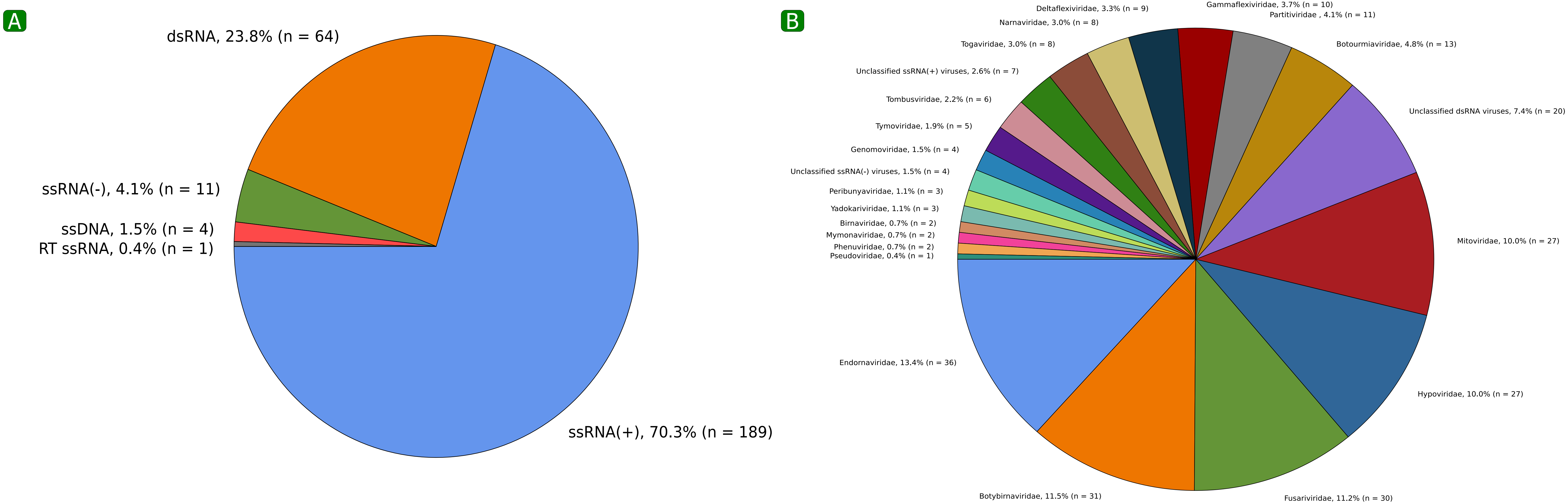
A
dsRNA, 23.8% (n = 64)
ssRNA(-), 4.1% (n = 11)
ssDNA, 1.5% (n = 4)
RT ssRNA, 0.4% (n = 1)
ssRNA(+), 70.3% (n = 189)
B
Deltaflexiviridae, 3.3% (n = 9)
Gammaflexiviridae, 3.7% (n = 10)
Partitiviridae , 4.1% (n = 11)
Narnaviridae, 3.0% (n = 8)
Togaviridae, 3.0% (n = 8)
Botourmiaviridae, 4.8% (n = 13)
Unclassified ssRNA(+) viruses, 2.6% (n = 7)
Tombusviridae, 2.2% (n = 6)
Unclassified dsRNA viruses, 7.4% (n = 20)
Tymoviridae, 1.9% (n = 5)
Genomoviridae, 1.5% (n = 4)
Unclassified ssRNA(-) viruses, 1.5% (n = 4)
Peribunyaviridae, 1.1% (n = 3)
Yadokariviridae, 1.1% (n = 3)
Birnaviridae, 0.7% (n = 2)
Mitoviridae, 10.0% (n = 27)
Mymonaviridae, 0.7% (n = 2)
Phenuviridae, 0.7% (n = 2)
Pseudoviridae, 0.4% (n = 1)
Endornaviridae, 13.4% (n = 36)
Hypoviridae, 10.0% (n = 27)
Botybirnaviridae, 11.5% (n = 31)
Fusariviridae, 11.2% (n = 30)

**Fig 1** Phylogenetic trees of novel RNA-dependent RNA polymerase (RdRp) sequences in the **(A)** *Pisuviricota* phylum (n=48), **(B)** *Lenarviricota* phylum (n=33), **(C)** *Kitrinoviricota* phylum (n=38), **(D)** *Negarnaviricota* phylum (n=5), **(E)** family *Botybirnaviridae* (n=24), and **(F)** viruses unclassified at the phylum level (n=17) prior to performing any conserved protein motif analysis. Accession numbers in red are from this study (indicated in green in the outer ring). The RdRp-Scan whole tree backbone, or NCBI records were used to create the phylogenetic trees.

**Fig 2** Multiple sequence alignment (MSA) visualizations of RdRp conserved protein motifs (A, B, and C) of novel RdRps in the **(A)** *Lenarviricota* phylum (n=17), **(B)** *Kitrinoviricota* phylum, (n=17), **(C)** in the *Pisuviricota* phylum (n=16), and **(D)** *Negarnaviricota* (n=4) using MegAlign Pro v. 17.5.0 (DNAStar Lasergene software).

**Fig 3** Multiple sequence alignment (MSA) visualizations of **(A)** Replication initiation protein conserved protein motifs (A, B, and C) of novel contigs in the *Genomoviridae* family (n=1), of **(B)** RNA-dependent RNA polymerase (RdRp) conserved protein motifs (A, B, and C) of novel contigs in the family *Botybirnaviridae* (n=5), of **(C)** RdRp conserved protein Motifs A-E of novel contigs in the family *Yadokariviridae* (n=2), and of **(D)** RdRp conserved protein Motifs A-E of novel contigs related to Botrytis cinerea mycovirus 5 and related dsRNA mycoviruses (n=1) using MegAlign Pro v. 17.5.0 (DNAStar Lasergene software).

**Fig 4** Multiple sequence alignment (MSA) visualizations of RNA-dependent RNA polymerase (RdRp) conserved protein motifs of novel contigs with **(A)** motifs I-VIII in the family *Quadriviridae* (n=2), **(B)** motifs A-D related to 'Orfanplasmoviruses' (n=2) and **(C)** motifs A-E related to Sclerotinia sclerotiorum dsRNA mycovirus L and related dsRNA mycoviruses (n=1) using MegAlign Pro v. 17.5.0 (DNAStar Lasergene software).

**Fig 5** Mycovirus genome types, families, and species identified in 44 isolates of Botrytis cinerea. **(A)** Percentage of mycoviruses belonging to ssRNA(+), ssRNA(-), dsRNA, ssDNA and RT ssRNA genome types identified. **(B)** Percentage of mycoviruses belonging to different families identified.
